# kitAb: an automated framework for assay-specific antibody developability modeling and large-scale landscape analysis

**DOI:** 10.64898/2026.10.09.750540

**Authors:** Ekaterina Trofimova, Elizaveta Shaburova, Denis Antonets, Vasily Ramensky

## Abstract

Antibody developability is determined by multiple physicochemical properties, and the relevance of individual properties varies among experimental assays. Existing computational approaches, however, often provide fixed assessments or rely on computationally intensive descriptors. We present kitAb, an open-source framework for scalable physicochemical characterization of antibody Fv regions and assay-specific developability modeling. kitAb calculates 59 interpretable sequence- and structure-based descriptors and integrates them with automated feature selection and regression.

We evaluated kitAb across ten public datasets comprising 46 experimental endpoints, using sequence-aware or predefined splits where feasible. Predictive performance was strongly assay-dependent. In nested evaluations with sequence-aware or predefined data splits, models using kitAb descriptors achieved predictive performance comparable on average to those using PROPERMAB descriptors, while outperforming models based on TAP or sequence-only features. In a single-structure CPU timing benchmark, kitAb descriptor calculation was approximately 11-fold faster than PROPERMAB. Predictive performance was similar across the three structure-generation backends, and broader AutoML searches did not substantially improve mean performance.

We further applied the same descriptor representation to approximately 3.4 million paired natural antibodies and 706 Phase II or later therapeutic antibodies. Repertoire-scale analysis revealed systematic differences in surface charge, hydrophobic and aromatic organization, CDR length and germline-associated physicochemical profiles. The kitAb code and analysis workflows are openly available at https://github.com/katyachemistry/kitAb/.

## Introduction

Developability describes whether an antibody with the desired binding properties can be manufactured and advanced as a therapeutic. It encompasses properties such as expression yield, aggregation propensity, colloidal stability, high-concentration viscosity, and nonspecific interactions [1]. Because liabilities may emerge late, when correction is costly and otherwise promising candidates may be deprioritized, they should be assessed early in discovery and optimization.

Computational developability assessment aims to triage candidates before experimental characterization. Many approaches use interpretable physicochemical descriptors, including exposed hydrophobicity, net charge, charge asymmetry, and electrostatic or hydrophobic surface patches. Spatial Aggregation Propensity (SAP) [2] identifies aggregation-prone regions formed by spatially adjacent hydrophobic residues, the Developability Index [3] combines SAP with net charge, and Spatial Charge Map (SCM) [4] quantifies negatively charged Fv surface environments associated with high-concentration viscosity. Surface curvature has also been shown to be predictive of antibody aggregation [5]. General protein solubility and aggregation predictors are also applicable to antibody assessment.

Deep-learning approaches to antibody developability are constrained by the scarcity of public datasets, which usually contain only tens to hundreds of measurements, are assay-specific, and are difficult to compare directly. Nevertheless, sequence-based models have been used to approximate computationally expensive spatial descriptors, as in DeepSCM [6], and to predict experimental developability measurements in low-data settings [7]. Reference-based methods instead assess how strongly candidate properties deviate from distributions observed for therapeutic antibodies; the Therapeutic Antibody Profiler (TAP) formalized this approach using interpretable structure-based descriptors [8]. More recent approaches use protein language models (pLMs) for zero-shot scoring or fine-tuned prediction, and increasingly combine pLM representations with molecular descriptors [9,10].

Despite this progress, infrastructure for scalable antibody developability assessment remains fragmented. TAP offers web-based assessment with a compact descriptor set but is not distributed as a standalone public codebase. Molecular Operating Environment (MOE) provides a comprehensive set of molecular descriptors, but is a commercial product. PROPERMAB [11] is open source, although its use of computationally intensive electrostatic potential and molecular surface calculations can increase runtime for large libraries. Thus, the existing tools typically provide either descriptor-based assessments or endpoint-specific predictors, rather than a single fast, open-source workflow that combines scalable descriptor calculation with assay-specific model development.

Public developability datasets remain small and cover only a limited range of antibodies, assays, and experimental conditions [12]. Researchers can generate developability measurements for their own candidates, but such experiments are costly and are therefore usually performed on relatively small sets of antibodies. This creates a need for assay-specific predictive models that can learn from limited experimental data and then be applied to prioritize much larger candidate libraries.

Here, we present kitAb, an open-source toolkit for scalable physicochemical characterization of antibody Fv regions and assay-specific developability modeling. The toolkit integrates structure prediction, physicochemical descriptor calculation, and automated model training and comparison. We benchmark kitAb across public developability datasets and apply the same descriptor panel to millions of predicted Fv structures. Rather than defining a universal developability score, kitAb provides a representation based on common physicochemical features that can be used either to learn assay-specific predictors from labeled data or to characterize physicochemical variation across large antibody collections.

## Methods

Detailed software settings, residue-class definitions, minimization parameters, descriptor formulas, and analysis definitions are provided in the Supplementary Section 1.

### Data preparation

Therapeutic antibody sequences were downloaded from Thera-SAbDab [13] on 21 June 2026. We removed records classified as TBC, Unknown, Phase I, Phase I/II, Preclinical, or withdrawn, as well as entries lacking usable paired heavy and light chain sequences. Duplicate concatenated heavy-light sequences were collapsed, yielding 706 unique paired Fv sequences spanning Phase II or later development stages, including approved antibodies. V- and J-gene assignments were obtained with ANARCI using IMGT numbering and germline assignment.

Natural paired human antibodies were assembled from 3,003,127 OAS [14] (accession date 7 May 2025) and 4,930,995 pre-filtered PairedAbNGS sequences [15] (accession date 1 July 2025). CDR and framework regions were defined using IMGT annotation [16]; PairedAbNGS sequences were annotated and assigned germline genes with ANARCI [17]. We excluded selected disease-associated repertoires, incomplete or invalid annotations, mouse-germline artefacts [18], and highly redundant sequences identified by MMseqs2 clustering of concatenated CDRs at 95% identity. Missing N-terminal FW1 residues were restored from the corresponding germline genes. Structure preparation and minimization after ABodyBuilder2 (ABB2) prediction failed for 167,446 of 3,605,625 antibodies (4.64%), leaving 3,438,179 antibodies with successfully computed descriptors. To assess potential bias in the germline composition of the retained cohort, germline usage was compared between antibodies with and without successfully computed descriptors using two-sided Fisher’s exact tests with Benjamini-Hochberg correction; detailed results are provided in Supplementary Section 1.

Datasets for assessing AutoML performance were obtained from the sources listed in Table 1. In the context of the AutoML analyses, the terms “endpoint” and “target” are used interchangeably to refer to the experimentally measured assay outcome being predicted. Antibody sequences and target values were retained as reported in the sources, except that the computationally derived Fab pI target was excluded from the Jain 2024 dataset [19].

**Table 1.** Public datasets used for benchmarking kitAb on developability endpoint prediction.

| Dataset | Size | Classification | Targets | Properties |
| --- | --- | --- | --- | --- |
| AB21 [36] | 21 | Diverse | viscosity | Viscosity |
| Garbinski | 86 | Several parents | Tm1 | Thermostability |
| Ginkgo [37] | 93-2<br>41 | Diverse | SEC Monomer, SMAC, HIC, HAC, PR CHO, PR Ova, AC-SINS pH6.0, AC-SINS pH7.4, Tonset, Tm1, Tm2 | Thermostability<br>Aggregation<br>Polyreactivity |
| Hutchinson [38] | 192 | Single parent | Tm1 | Thermostability |
| Jain 2017 [39] | 137 | Diverse | AC SINS, BVP ELISA, CIC RT, CSI BLI, ELISA, HIC RT, PSR, SAS, SGAC SINS, SMAC RT, Tm | Thermostability<br>Aggregation<br>Polyreactivity |
| Jain 2023 [40] | 112-115 | Diverse | DNP, FA2, FVC32, FvFVIII2, FvLysM2, Heme2, HemeC32, HemeFVIII2, HemeLysM2 | Polyreactivity |
| Jain 2024 [19] | 43 | Diverse | AC-SINS, CIC RT, CS-SINS, HIC RT, Heparin RT, PSR, SEC, Tm, cIEF | Thermostability<br>Aggregation<br>Polyreactivity |
| Jetha 2019 [41] | 97 | Single parent | HIC RT | Aggregation |
| Kraft [42] | 128 | Diverse | Heparin RT | Aggregation<br>Polyreactivity |
| PDGF38 [43] | 38 | Single parent | viscosity | Viscosity |

Antibody Fv structures were predicted using ABodyBuilder2, ABodyBuilder3 (ABB3), and FlashABB [20–22], with inference on an NVIDIA A100 80 GB GPU and each method’s default pretrained weights. To assess cross-method variability and within-method reproducibility, each predicted structure underwent three independent structure-preparation and restrained-minimization runs. As detailed in Supplementary Section 1, missing atoms and hydrogens were added with PDBFixer and OpenMM and structures were refined with an ABB2-adapted restrained minimization protocol using the AMBER ff14SB force field.

### Descriptors

The framework comprised established and custom developability-related descriptors calculated from the Fv sequence and predicted structure, either over the full Fv or within the CDR vicinity, defined as all IMGT CDR residues plus residues with at least one heavy atom within 5 Å of any CDR heavy atom. Residues were grouped into negatively charged, positively charged, aromatic, and hydrophobic classes; charge class assignments used PROPKA-derived fractional charges at pH 7.5. Residues with relative side-chain solvent-accessible surface area (SASA) ≥20% were considered exposed. Full residue-class definitions, formulas and descriptor names are provided in Supplementary Section 1.

### Surface physicochemistry and spatial organization

Residue-level SASA values were calculated with FreeSASA 2.1.2 using the Shrake-Rupley algorithm and Ala-X-Ala reference values [23]. Two sets of SASA descriptors were calculated: global descriptors spanning the entire Fv and local descriptors restricted to the CDR vicinity. For each aromatic, charged, and hydrophobic residue class, global descriptors included total side-chain SASA and the fraction of this SASA contributed by solvent-exposed residues. CDR-vicinity descriptors included an analogous measure of the side-chain SASA of solvent-exposed residues, together with an additional measure of their average relative exposure.

Residue hydrophobicity was assigned using the Wimley-White interfacial scale [24]. Global hydrophobicity was calculated by summing residue-level hydrophobicity values across the Fv. CDR-vicinity hydrophobicity was calculated as an exposure-weighted mean, giving greater weight to more solvent-exposed residues.

Spatial patches were identified globally and within the CDR vicinity using DBSCAN on Cα coordinates. All residues of each class were included, with charged residues clustered using ε = 10 Å and min_samples = 2 and hydrophobic residues using ε = 5 Å and min_samples = 3. Noise points were excluded, and patch area was defined as the summed side-chain SASA of clustered residues. For positively and negatively charged residues, Shannon entropy of cluster-level SASA described whether charged surface area was concentrated in one patch or distributed across several patches.

Pair-correlation descriptors measured enrichment of unordered same-class Cα pairs of exposed negatively charged, positively charged, and hydrophobic residues within 4-7 Å and 7-10 Å shells. Pair counts were normalized against references generated by repeatedly sampling the same number of residues from all exposed Cα positions. Values above or below 1 indicated enrichment or depletion; classes with fewer than two residues were assigned 0.

### Local geometry and hydrogen bonding

Local planarity descriptors characterized the geometric flatness of exposed CDR-vicinity residues from each physicochemical class. For each heavy atom, neighboring heavy atoms within 8 Å were used to compute the 3D covariance of their coordinates, and planarity was defined as the fraction of variance along the thinnest principal axis (smallest eigenvalue divided by the sum of all three eigenvalues), with lower values indicating a flatter local arrangement. Atom-level values were averaged within residues; class-specific residue scores were then summed and normalized by the mean score across all exposed CDR-vicinity residues (Supplementary Section 1).

Two complementary H-bond descriptors were calculated. CDR H-bond density used a custom geometry-based detector that included backbone and side-chain H-bonds, with donor and acceptor contributions weighted by relative main- or side-chain SASA. H-bond energy density used DSSP-annotated backbone H-bonds (DSSP 4.6.1 [25]), with absolute interaction energies weighted by the participating residues’ relative main-chain SASA. Full geometric criteria and normalization are provided in Supplementary Section 1.

### Global and sequence-derived properties

Unless stated otherwise, charge descriptors were calculated using PROPKA-derived fractional residue charges at pH 7.5. Global properties included net charge, isoelectric point, heavy-light charge symmetry, dipole moment calculated from AMBER ff19SB partial atomic charges relative to the atomic centroid, and heavy-light interface burial measured as SASA lost upon chain association. PROPKA 3.5.1 [26] was used to estimate residue protonation states for charge-related descriptors. ff19SB partial atomic charges were also used for the SCM-like spatial charge descriptors and were treated separately from the PROPKA-derived residue-level charges (Supplementary Section 1).

CDR composition descriptors quantified glycine, proline, aromatic, positively charged, negatively charged, and glutamine/asparagine fractions within the strict CDR set. Asp-Asp, Asp-Glu, and Tyr-Tyr motifs were counted in CDR-vicinity sequence fragments. Salt bridges were defined as unique oppositely charged residue pairs with at least one interatomic contact within 4 Å.

### AutoML and benchmarking

The AutoML pipeline was run separately for each dataset and endpoint (Table 1). Where feasible, MMseqs2 [27] was used to generate sequence-aware folds at 0.80 sequence identity; Garbinski, Kraft, Jain 2017, Jain 2023, and Jain 2024 were divided into four or five balanced folds. Ginkgo predefined folds were retained, whereas Jetha, Hutchinson, AB21, and PDGF38 used random five-fold splits repeated across three seeds because sequence-aware partitioning was infeasible. Four modeling strategies were evaluated: linear-kernel support vector regression (SVR) after low-variance and correlation filtering; sequential forward feature selection guided by either linear-kernel SVR or k-nearest-neighbour regression; and elastic-net regression without feature selection. For datasets with >80 samples, model strategy and internal settings were selected by inner cross-validation within each outer training fold, then refit on the full outer training set and evaluated on the held-out fold. Datasets with <80 samples used flat CV. Feature usage was summarized for the strategy with the highest mean inner-CV Spearman across outer folds: ElasticNet and correlation-pruned SVR models were scored by mean L1-normalized absolute coefficients, and SFS models by selection frequency across outer folds. Assay-group enrichment was tested within each representation and modeling strategy using two-sided Mann-Whitney U tests with Benjamini-Hochberg correction.

Benchmark representations comprised TAP scores, 31 PROPERMAB sequence- and structure-based descriptors, and the PROPERMAB sequence-only subset (CDR-H3 length, CDR aromatic count, theoretical pI, number of charged Fv residues, Fv charge, and Fv charge symmetry). TAP scores were obtained directly from antibody sequences using FlashABB [22], whereas structural PROPERMAB descriptors were calculated from the same minimized Fv structures used for kitAb. For Ginkgo, externally provided heavy and light chain subtype annotations were encoded as indicator variables and appended identically to all four representations. Their contribution was assessed by repeating the Ginkgo ABB2 nested-CV analysis after removing these indicators while retaining the original folds and model-selection procedure.

Wall-clock descriptor-calculation times for kitAb and PROPERMAB were measured on an Intel Xeon Gold 6348 workstation using 50 single-threaded repeats on the same antibody structure (PDGF38 AB-001), with math libraries pinned to one thread. kitAb timings included DSSP, PropKa, FreeSASA, and descriptor calculation; PROPERMAB timings included its full preprocessing and molecular feature-generation workflow from the same PDB input. These values therefore represent a controlled single-structure comparison rather than hardware-independent absolute throughput.

## Results

### kitAb integrates antibody characterization and developability prediction with reproducible structure-based descriptors

The kitAb pipeline was designed to provide computationally inexpensive and interpretable descriptor calculation for characterizing developability-related physicochemical properties at library scale and for predicting assay outcomes from labeled data in a fully automated manner. Rather than relying on fine-grained characterization through computationally intensive molecular simulations, surface modeling, or electrostatic calculations, kitAb derives a fixed set of aggregate, coarse-grained descriptors from static Fv structure and sequence, as detailed in the Methods and Supplementary.

The user-configurable workflow comprises three stages: optional Fv structure prediction and minimization, calculation of standardized global and local descriptors, and optional assay-specific AutoML model development (Fig. 1). To use the AutoML module, the user provides a labeled training dataset containing antibody Fv sequences or structures, corresponding experimental measurements for the developability endpoints of interest and, optionally, external features and predefined cross-validation folds. kitAb then assesses feature selection and regression strategies and trains an endpoint-specific predictive model.

**Figure 1.**
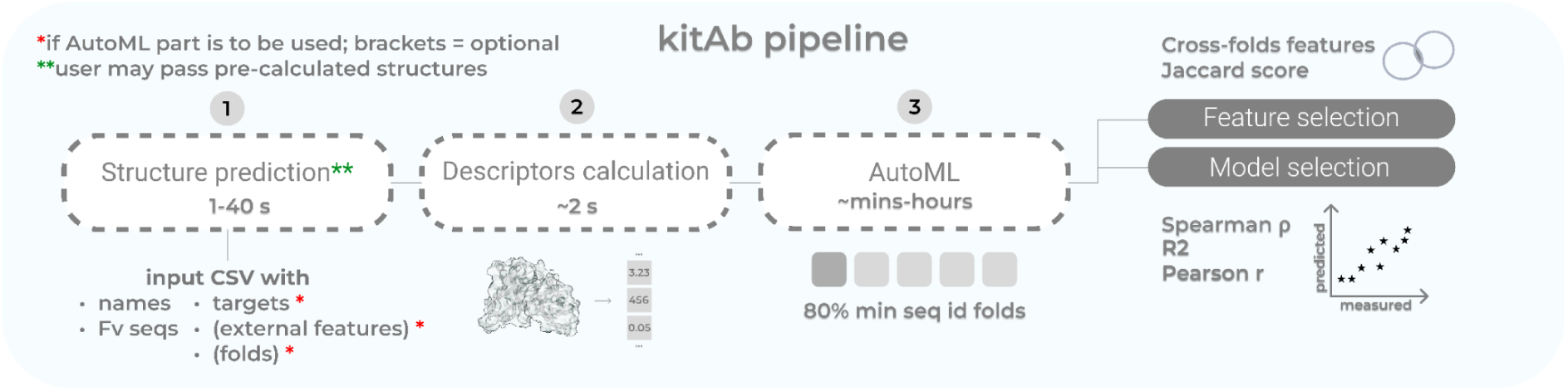
Overview of the kitAb workflow. Optional Fv structure prediction and minimization are followed by descriptor calculation and, when labeled data are provided, optional AutoML with nested or flat CV.

The kitAb panel comprises 59 computationally efficient descriptors spanning surface composition, spatial organization, intramolecular interactions, and global physicochemical properties. SASA-based descriptors quantify hydrophobic, aromatic, and charged surface composition through absolute and relative side-chain exposure, whereas planarity descriptors summarize the local geometric organization of residues. Hydrophobicity-weighted absolute SASA has previously been associated with hydrophobic interaction chromatography (HIC) retention time [28]; accordingly, kitAb uses the relative-SASA-weighted hydrophobicity score, based on the Wimley-White scale, which performed well in prior comparisons against antibody HIC measurements [24,28]. kitAb also introduces three lightweight SAP-inspired [2] descriptors capturing exposed aromatic surface and local aromatic-charge co-occurrence [29–31] and a positive-charge analogue of the viscosity-associated SCM descriptor [4]. Patch area descriptors reflect both accessible hydrophobic or charged surface and its clustering, while cluster entropy distinguishes concentrated from distributed charged patches [31]. Pair-correlation enrichment complements these measures by quantifying whether exposed residues of the same class occur together more often than expected by chance. The panel also includes thermostability-related features: salt bridges, CDR-vicinity hydrogen bonding, DSSP-derived backbone H-bond energetics, and heavy-light interface burial [32–34], as well as commonly used descriptors such as pI, CDR length, net charge, and charge symmetry. CDR composition is summarized by the fractions of Gly, Pro, aromatic, positively charged, negatively charged, and Gln/Asn residues, together with Asp-Asp, Asp-Glu, and Tyr-Tyr motif counts [29,35]. Overall, kitAb combines established and custom descriptors selected for computational efficiency and mechanistic interpretability.

We assessed descriptor sensitivity to the structure-prediction method and repeated processing using the non-redundant set of antibodies represented across the public datasets used later for model development (Table 1). Between-method normalized variability ranged from approximately 0.1 to 1.3 (median 0.35; Fig. 2, Fig. S1) and was highest for H-bond and PCF descriptors. Descriptors with normalized ranges above 0.5 also included planarity scores, inter-chain buried SASA, salt-bridge counts, and exposed-SASA features. These descriptors depend strongly on local geometry or exposure thresholds, making them sensitive to small structural shifts that alter detected hydrogen bonds or salt bridges, or change which residues satisfy the exposure criterion.

**Figure 2.**
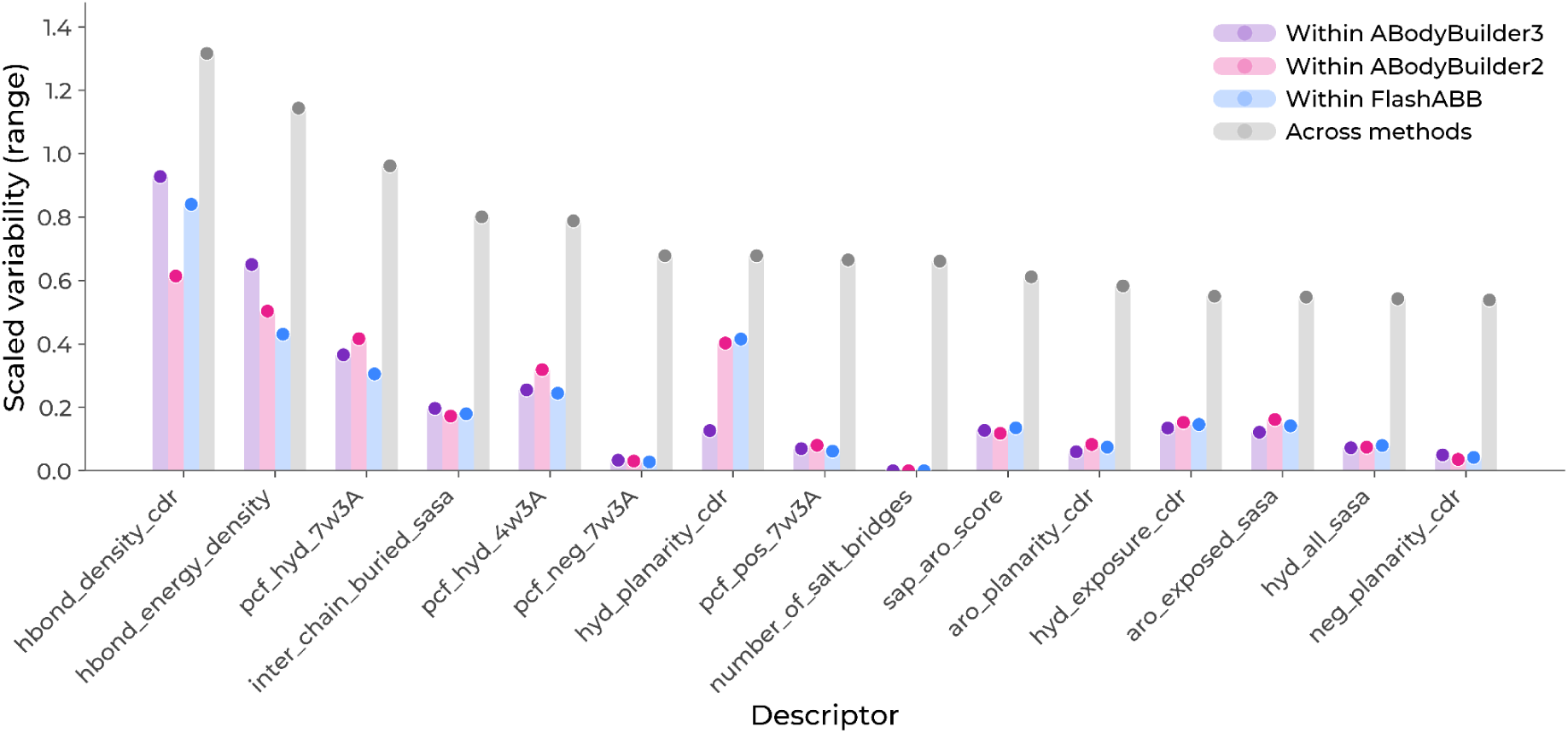
Fifteen descriptors with the highest variability across structure-prediction methods. Variability is expressed as the across-method descriptor range normalized by between-antibody variation (see Supplementary Section 1 for details).

Within-method variability across repeated minimization runs was much lower (median 0.07-0.08), with no clear method-specific bias. ABB2 predictions were somewhat more divergent from the other two structure backends: in 59% of antibody-descriptor combinations, the difference between ABB2 and at least one other method exceeded the corresponding ABB3-FlashABB difference.

### kitAb provides competitive and interpretable predictive performance at reduced computational cost

We evaluated the kitAb AutoML pipeline across ten public antibody developability datasets spanning 46 endpoints related to thermostability, aggregation, polyreactivity, and viscosity (Table 1). The datasets contained 21-241 antibodies and included both diverse antibody panels and shared-parent variant series. For sufficiently large endpoints (>80 samples), nested cross-validation was used to estimate performance after AutoML selection; smaller datasets were evaluated without nesting. The benchmark used the common Fv-centered representation employed in many antibody developability studies and thus primarily assessed Fv-associated predictive signal.

Among endpoints evaluated by nested cross-validation with sequence-aware or predefined outer folds (n = 33), kitAb and PROPERMAB performed similarly, with Fisher-z-mean Spearman ρ = 0.36, compared with 0.27 for the sequence baseline and 0.18 for TAP (Fig. 3A). For the smaller Jain 2024 dataset, which was evaluated by non-nested flat CV, mean ρ was 0.43 for kitAb, 0.36 for PROPERMAB and the sequence baseline, and 0.21 for TAP. Random splitting was used primarily for shared-parent variant series and reflects local interpolation within an engineered sequence neighborhood rather than across-antibody generalization. In these random-split evaluations, mean ρ was 0.54 for kitAb, 0.58 for PROPERMAB, 0.42 for the sequence baseline, and 0.28 for TAP.

**Figure 3.**
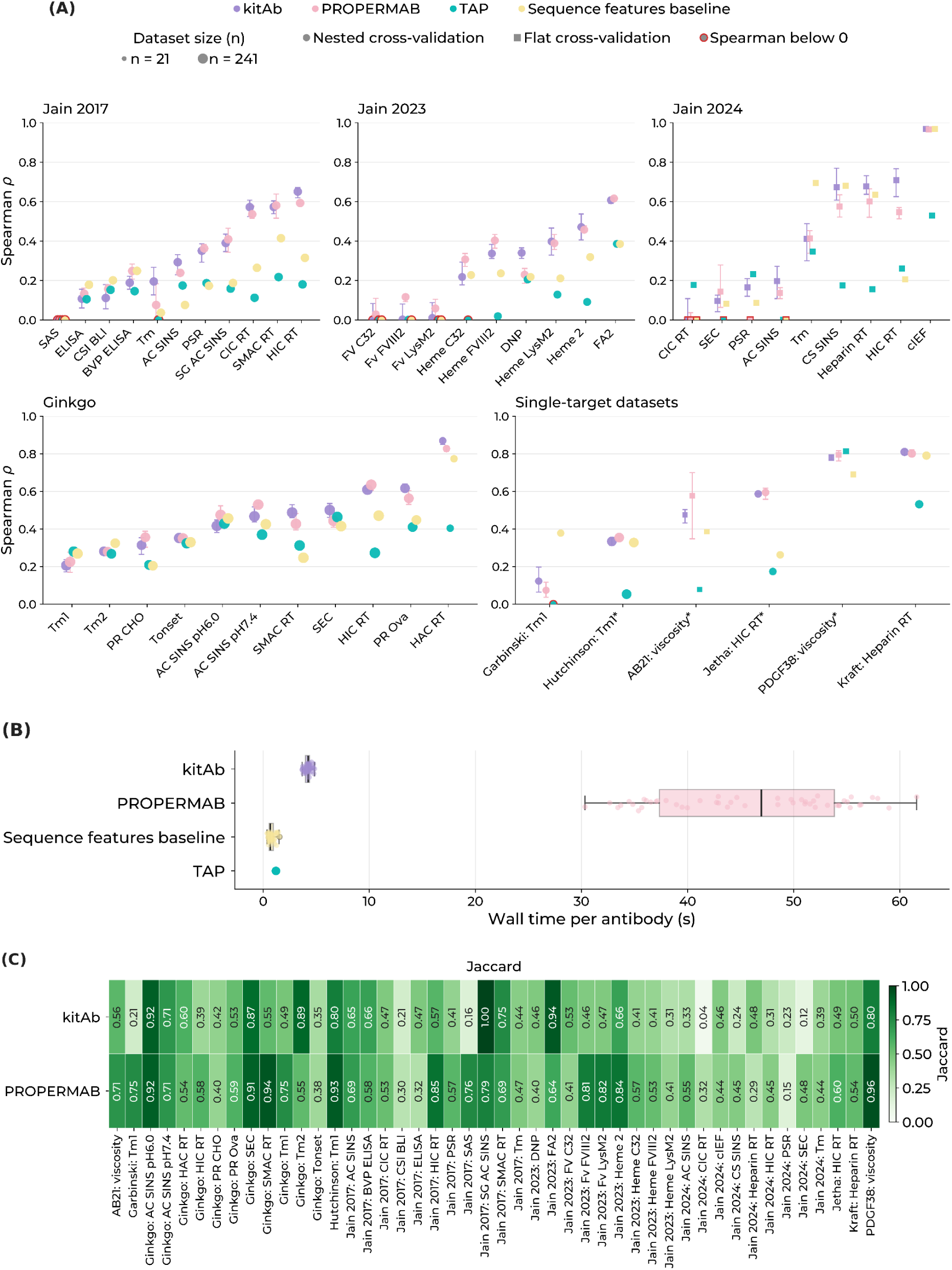
(A) Assay-level averaged Spearman *ρ* from pooled out-of-fold (OOF) predictions. Fisher-z means were calculated across repeated processing runs and then across structure-generation backends; error bars show the min-max range across backends. An asterisk denotes random-split evaluations. (B) Descriptors calculation time benchmark. (C) Feature recurrence for one modeling strategy per run, selected by its mean inner-CV performance across outer folds; mean pairwise Jaccard agreement was then calculated among the feature sets selected in the outer folds.

Repeated structure-processing runs for the same backend yielded similar kitAb predictive performance. Run-to-run differences in performance were not associated with descriptor variability. They were also unrelated to feature-set overlap across outer folds (ABB2, ABB3, and FlashABB: Spearman ρ = −0.07, −0.07, and 0.01; all p > 0.6). In mixed-effects analysis, assay group showed the strongest association with predictive performance (p = 9.4 × 10⁻⁷), whereas structure-generation backend and split strategy had no detectable effect. Effective sample size and CV nestedness showed nominal associations, but these could not be cleanly separated from dataset identity; the assay-group effect remained significant when dataset identity was included. No assay group showed a significant preference for a particular modeling strategy after FDR correction in either kitAb or PROPERMAB. For kitAb, ElasticNet was selected most often (52% of outer folds). Feature-set stability, however, varied substantially between endpoints (Fig. 3C), with recurrence ranging from 0.04 to 1.00 for kitAb and from 0.15 to 0.96 for PROPERMAB.

We next asked whether a more extensive AutoML search could improve performance. Across 35 endpoints evaluated by nested CV, we compared the standard kitAb pipeline with a broader search (correlation and stability filtering, recursive feature elimination and SFS, multiple feature budgets, and linear and nonlinear models) and with an established AutoML tool, AutoGluon Tabular [44]. This comparison used a single ABB2 structure-generation run for each endpoint. Fisher-z mean Spearman ρ was 0.38, 0.39, and 0.28 for the standard pipeline, broader search, and AutoGluon, respectively. Thus, expanding the search space did not significantly improve mean predictive performance despite increasing runtime by approximately an order of magnitude, and AutoGluon yielded a lower mean correlation. We therefore retained the standard kitAb AutoML configuration for subsequent analyses.

Predictive accuracy varied by assay type, with hydrophobicity-related endpoints among the most consistent (Fig. 3A). For HIC, kitAb achieved Spearman ρ = 0.59-0.71 across Jetha, Ginkgo, Jain 2017, and Jain 2024, approaching the MOE baseline of 0.66 reported for Ginkgo [45] (kitAb ρ = 0.61) and outperforming TAP and the sequence-only baseline, consistent with prior evidence that chromatographic hydrophobicity is accessible to structural physicochemical descriptors. Stand-up monolayer adsorption chromatography (SMAC) was also predicted well (ρ = 0.49-0.57), whereas salt-gradient affinity-capture self-interaction nanoparticle spectroscopy (SG AC-SINS) reached 0.39. Feature usage supported a coherent hydrophobicity signature. Individual HIC and SMAC models emphasized hydrophobic exposure, hydrophobic patches, and aromatic surface descriptors, including Tyr-Tyr content. CDR hydrophobic exposure was the strongest hydrophobicity contrast among kitAb coefficient models (Δ = 5.53 percentage points) but was not significant after FDR control. In contrast, SG AC-SINS combined hydrophobic and aromatic descriptors with charge symmetry and dipole moment, suggesting a broader electrostatic contribution.

Self-association was less uniformly captured than hydrophobicity-related behavior, with predictability depending strongly on the assay. AC-SINS was moderate for Ginkgo at pH 6.0 and 7.4 (ρ = 0.42 and 0.47) and weaker for Jain 2017 (ρ = 0.29). Because the Ginkgo benchmark included externally provided chain subtype annotations, we assessed their contribution by repeating the nested-CV analysis (ABB2, single run) after removing these indicators from all compared representations (Fig. S9). Across the 11 Ginkgo endpoints, removal of subtype features reduced mean Spearman ρ by 0.15 for kitAb, 0.13 for PROPERMAB, 0.20 for TAP, and 0.18 for the sequence baseline, with the largest declines observed for stability endpoints, but also for AC-SINS. Clone self-interaction by biolayer interferometry (CSI-BLI) was poorly predicted; charge-stabilized SINS (CS-SINS) reached ρ = 0.67 and was matched by the sequence baseline, driven mainly by pI and charged-residue features. Across AC-SINS endpoints, recurrent physicochemical features included dipole moment, SCM charge descriptors, Gln and Asn content, and hydrophobic and aromatic surface descriptors, broadly consistent with previous findings implicating both electrostatic and hydrophobic properties in self-association [46].

Cross-interaction behavior was highly dataset-dependent. kitAb predicted cross-interaction chromatography (CIC) well in Jain 2017 (ρ = 0.57), but not in the smaller Jain 2024 dataset; similarly, polyspecificity reagent (PSR) binding was only moderate in Jain 2017 (ρ = 0.35) and weak in Jain 2024. No feature enrichment survived FDR correction, but exposure-weighted CDR hydrophobicity showed the largest shift. The underlying feature patterns were interpretable: Jain 2017 CIC emphasized hydrophobicity and aromatic surface properties, including Tyr-Tyr content, whereas PSR showed an electrostatic signature involving positive SCM, dipole moment, charge symmetry and net charge. Negative-charge descriptors were also negatively correlated with both CIC and PSR outcomes (Fig. S4), indicating that greater negative-charge character was associated with lower nonspecific interaction propensity. These observations are consistent with evidence that CIC partly reflects hydrophobic interactions [47] and that CDR hydrophobicity and positive charge can favor polyreactivity, whereas acidic CDR residues have been associated with greater specificity [48]. PROPERMAB showed a similar overall profile.

Polyreactivity endpoints varied substantially in predictability. kitAb performed best for polyreactivity to ovalbumin (PR Ova; ρ = 0.62), while 2,4-dinitrophenol (DNP) binding and polyreactivity to CHO cells (PR CHO) were intermediate, and baculovirus particle ELISA (BVP ELISA) and the six-antigen ELISA remained weak. Feature usage pointed mainly to surface-charge organization, including negative-cluster entropy, SCM positive charge, and positive CDR charge exposure. The sequence baseline was lower than kitAb for PR Ova (ρ = 0.45) but performed better on all ELISA endpoints.

No kitAb descriptor was significantly enriched for viscosity. PDGF38 remained among the best-predicted targets for kitAb (ρ = 0.78), although the models using compact TAP descriptor set performed slightly better (ρ = 0.81), suggesting that a small set of surface properties was sufficient for this series. Those TAP-based models relied primarily on positive-charge and hydrophobicity descriptors. AB21 was more difficult; in random-split evaluation, PROPERMAB exceeded kitAb (ρ = 0.58 vs. 0.48) and emphasized negative patches and other charge-related descriptors.

Thermostability endpoints remained heterogeneous and generally difficult to predict. In Ginkgo, subtype annotations were frequently selected for several stability-related endpoints; notably, IgG4 heavy chain subtype was selected in every fold for Tm1 [49], consistent with the strong sensitivity of Tm1, Tm2, and Tonset to subtype removal (Fig. S9). Among the physicochemical descriptors, heavy-light inter-chain burial contributed to Ginkgo Tm2 and hydrophobic surface area to Tonset. Jain 2024 Tm emphasized CDR proline content, hydrophobicity, charge symmetry, salt bridges, and inter-chain buried SASA. Nevertheless, structural descriptors did not provide a consistent advantage for thermostability: the sequence baseline substantially outperformed kitAb for Jain 2024 Tm (ρ = 0.69 vs. 0.41) and Garbinski Tm1 (0.38 vs. 0.12).

Charge-related endpoints were among the most predictable. HAC and heparin retention were predicted well (ρ = 0.87 and 0.81), and cIEF was captured almost perfectly (ρ = 0.97). These endpoints were largely explained by simple electrostatic properties, as sequence-based descriptors nearly matched the structural models for cIEF and heparin retention. By contrast, FA2 (antibody binding to folate) and Heme2 (antibody binding to heme) showed a broader electrostatic signature involving dipole moment, charge symmetry, spatial charge organization, and charged surface patches.

Induced-polyreactivity assays measure binding acquired after exposure to Fe²⁺ or heme toward three unrelated proteins: complement component C3, coagulation factor VIII (FVIII), and the bacterial LysM domain. The heme-induced targets were consistently more predictable than their corresponding Fe²⁺ targets, reaching ρ = 0.22, 0.34, and 0.40 for C3, FVIII, and LysM, respectively. No kitAb coefficient-model enrichment survived FDR correction; aromatic all-SASA and exposed positive SASA showed the largest positive shifts (Δ = 6.84 and 3.17 percentage points; q = 0.069 and 0.052). Aromatic and exposed-surface descriptors repeatedly ranked highly in the heme-associated models.

Notably, the PROPERMAB sequence baseline outperformed full PROPERMAB for several endpoints by substantial margins. The largest differences were observed for Garbinski Tm1 (ρ = 0.38 vs. 0.07; Δρ = +0.31) and Jain 2024 Tm (ρ = 0.69 vs. 0.41; Δρ = +0.28), where sequence-only models also exceeded kitAb. For Jain 2024 PSR, the baseline outperformed PROPERMAB by Δρ = +0.21. Thus the larger descriptor space did not consistently translate into improved generalization, suggesting that in low-data settings, model and feature selection may fail to recover the most informative subset even when it is contained within the full representation.

Overall, kitAb predictive performance was similar across the three structure-generation backends and, in the sequence-aware or predefined evaluations, exceeded the lightweight TAP and sequence-only representations on average by Δρ = +0.18 and +0.09, respectively, while matching those of PROPERMAB. In the controlled single-CPU benchmark described above, kitAb reduced descriptor-calculation time from 45.98 to 4.21 s for the tested structure, corresponding to an approximately 11-fold speed-up relative to PROPERMAB. TAP and sequence descriptors were faster still (∼1 s per antibody), but performed worse on average. These results indicate that coarse-grained, interpretable physicochemical descriptors retain useful predictive signal across several developability assays while remaining compatible with library-scale analysis.

### The kitAb descriptor space enables repertoire-scale mapping of Fv physicochemical variation

Developability-related properties are typically considered when optimizing individual therapeutic candidates, but they can also be viewed as quantitative phenotypes of antibody sequence space. Their systematic characterization at repertoire scale has enabled efforts to map antibody biophysical landscapes [50]. The kitAb descriptor panel can be used to locate antibodies within a multidimensional Fv physicochemical space. We therefore applied the kitAb representation to approximately 3.4 million paired natural antibodies from Observed Antibody Space (OAS) and PairedAbNGS and to 706 Phase II or later therapeutic antibodies to examine how Fv physicochemical properties vary across repertoire subsets and germline backgrounds; data preprocessing is described in Methods.

For comparison across B-cell differentiation-state groups, we restricted this analysis to PBMC-derived natural antibodies, whereas the remaining repertoire-scale analyses used the full natural-antibody collection. IGHV3-23-containing heavy–light combinations showed distinct shifts across B-cell states and therapeutics. IGHV3-23 + IGKV3-20 remained common across all groups. IGHV3-23 + IGKV3-11 increased in memory and antibody-secreting populations and reached its highest frequency among therapeutics. IGHV3-23 + IGKV1-39 changed little across PBMC groups but was more frequent among therapeutics (2%). Several additional IGHV3-23-containing pairs also peaked in memory B cells. The prominence of some IGHV3-23-containing combinations may partly reflect the study composition of OAS, which contains substantial SARS-CoV-2 repertoire data; IGHV3-23/IGKV3-20, for example, was among the frequent heavy-light pairings reported in convalescent COVID-19 repertoires [51].

Because OAS and PairedAbNGS aggregate heterogeneous cohorts, biological sources, and experimental protocols, these patterns should be interpreted as cross-sectional differences among repertoire subsets rather than as maturation trajectories.

We next asked whether the repertoire differences identified above were accompanied by broader changes in the physicochemical properties of the antibody variable region (Fig. 4B). Several features differed across natural differentiation-state groups, and therapeutic antibodies did not simply resemble any one of these groups. Combined CDR-H3 and CDR-L3 length was highest in naïve antibodies, with a median of 25 residues compared with 22 in therapeutics. Longer CDR-H3 loops have been associated with polyreactivity in some studies, although this relationship is not consistent across datasets [52]. Therapeutic antibodies showed enrichment of proline in their CDRs (`fraction_pro_in_cdr` = 0.039 versus 0.021-0.022 across natural subsets).

**Figure 4.**
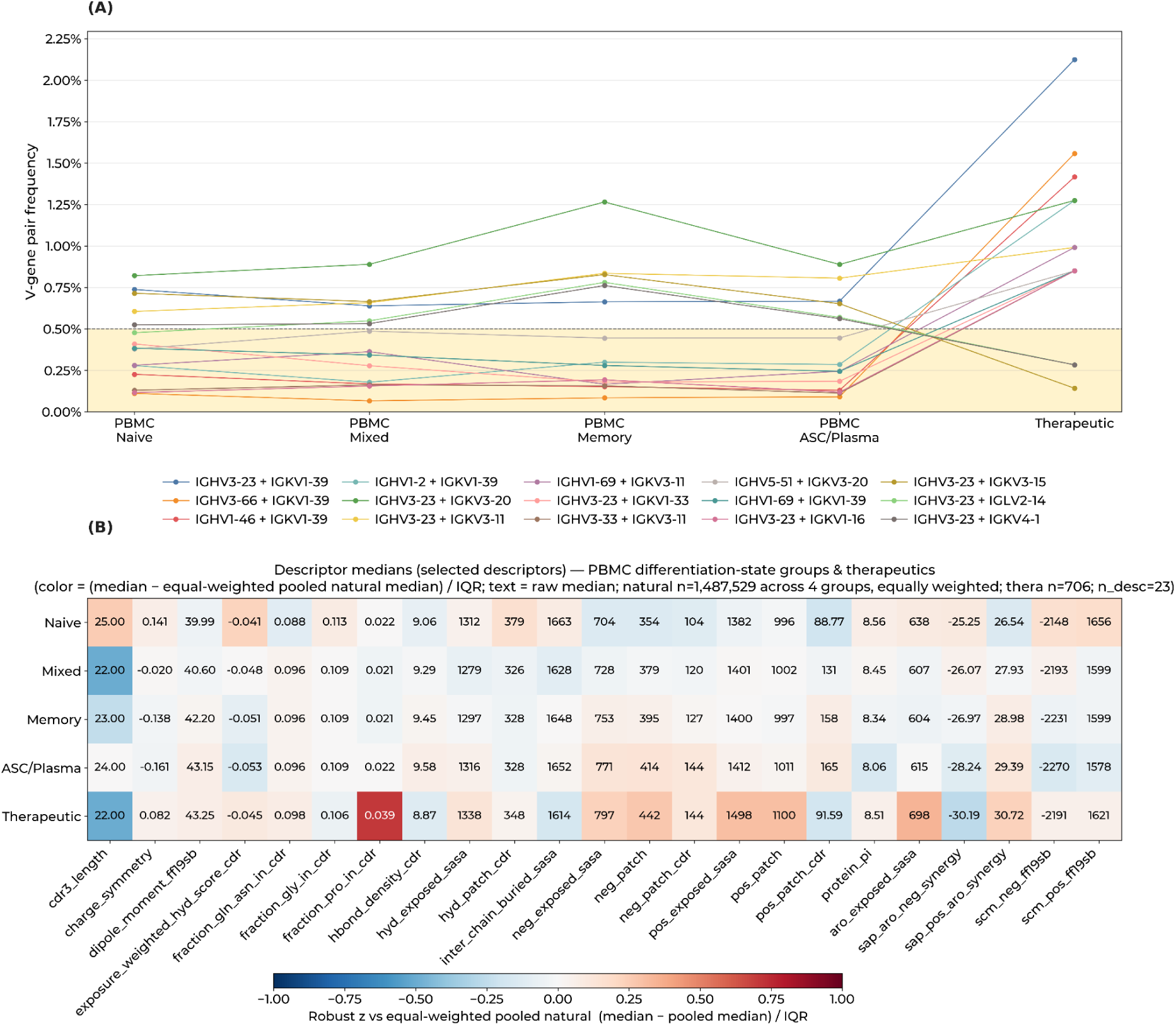
(A) Top 15 heavy-light V-gene pairs by maximum frequency among pairs with frequency ≥0.5% in PBMC naïve, Mixed, Memory, ASC/Plasma, or therapeutics. Frequencies are gene-level and normalized within each differentiation state group. (B) Median values of selected structural/physicochemical descriptors for PBMC B-cell differentiation states (naïve, Mixed, Memory, ASC/Plasma) and therapeutics. Color shows each group median as a robust z-score vs an equal-weighted pooled natural reference; text is the raw median.

Electrostatic organization showed a clear distinction between natural repertoire variation and the therapeutic set. Across the natural differentiation-state groups, both positive and negative CDR charge patches became larger as predicted pI decreased. This pattern suggests changes in the spatial organization of charge rather than a simple shift in net electrostatics. Therapeutics diverged from this pattern for positive charge: median `pos_patch_cdr` was close to the naïve value and markedly below the ASC/plasma value. Negative CDR patches behaved differently, with a median of 144 in both therapeutics and ASC/plasma antibodies. This asymmetry is notable because therapeutics were not globally depleted of positive charge; they instead had the largest overall positive exposed SASA and positive surface patches. Thus, the therapeutic set combined substantial global positive surface area with comparatively small positive patches around the antigen-binding region. Because concentrated positive charge near the paratope can promote nonspecific interactions, this pattern is consistent with preferential retention of therapeutic candidates lacking strongly positive CDR-localized patches.

Therapeutics had the highest exposed aromatic surface (`aro_exposed_sasa` = 698, versus 604-638 across the natural subsets), but this was not accompanied by an equally pronounced increase in CDR hydrophobic-patch area. The aromatic-charge synergy descriptors were strongest in therapeutics, suggesting that their exposed aromatic residues frequently occur in mixed charged-aromatic environments rather than in uniformly hydrophobic surface areas.

For repertoire-scale analyses, a 33-descriptor subset was selected from the full panel to define therapeutic-profile distance. Using this set, we tested whether germline genes were associated with greater or lesser distance from the multivariate therapeutic profile while accounting for biological source. Stratified rank effects ranged from -1 to +1, with negative values indicating greater similarity and positive values greater distance. The physicochemical representation and therapeutic-profile distance are also available through the web server https://kitab-atlas.com/ to support exploration and comparison of antibody profiles.

After requiring at least 2,000 antibodies, representation in ≥ 8 biological sources, a stringent heterogeneity filter (p ≥ 0.5), q ≤ 0.05, and an absolute rank effect ≥ 0.2 (see Supplementary Section 1 for analysis details), no heavy chain V genes passed the predefined robustness filters. The only light-chain V gene to meet the criteria, IGKV5-2, was strongly associated with greater distance from the therapeutic Fv profile (stratified rank effect 0.90). IGKV5-2 had a strongly acidic and electrostatically asymmetric profile, with increased negative-charge exposure and patching, reduced net charge and pI, and a markedly elevated dipole moment. IGKV5-2 has also been reported to be enriched among antibodies with low polyreactivity [53]. In that study, low-polyreactive sequences contained fewer positively charged residues and more aspartic acid, broadly consistent with the acidic profile observed here. This contrast illustrates the multidimensional nature of developability: association with a favorable individual property, such as low polyreactivity, does not necessarily imply overall similarity to the multivariate profile of therapeutic antibodies.

Heavy and light chain V-gene analysis identified multiple combinations associated with greater distance from the therapeutic profile. Among the strongest associations were IGHV4-31 × IGLV2-14, IGHV4-4 × IGLV2-14, IGHV4-59 × IGLV2-8, IGHV4-31 × IGLV1-40, IGHV4-34 × IGLV2-23, and IGHV3-9 × IGLV3-1, with stratified rank effects of 0.76-0.88. Five of these six combinations contained an IGHV4 gene, including four IGHV4 × IGLV2 pairings, and showed a shared pattern of altered surface charge organization relative to the therapeutic profile (Table 2). IGHV3-9 × IGLV3-1 differed from this pattern, with deviations concentrated in CDR-localized charge and hydrophobicity descriptors. Conversely, the three combinations most strongly associated with greater similarity to the therapeutic profile (stratified rank effect < -0.5; IGHV3-48 × IGKV1-12, IGHV3-23 × IGKV1-6, IGHV3-66 × IGKV1-39) all paired an IGHV3 heavy chain with an IGKV1 light chain.

**Table 2.**
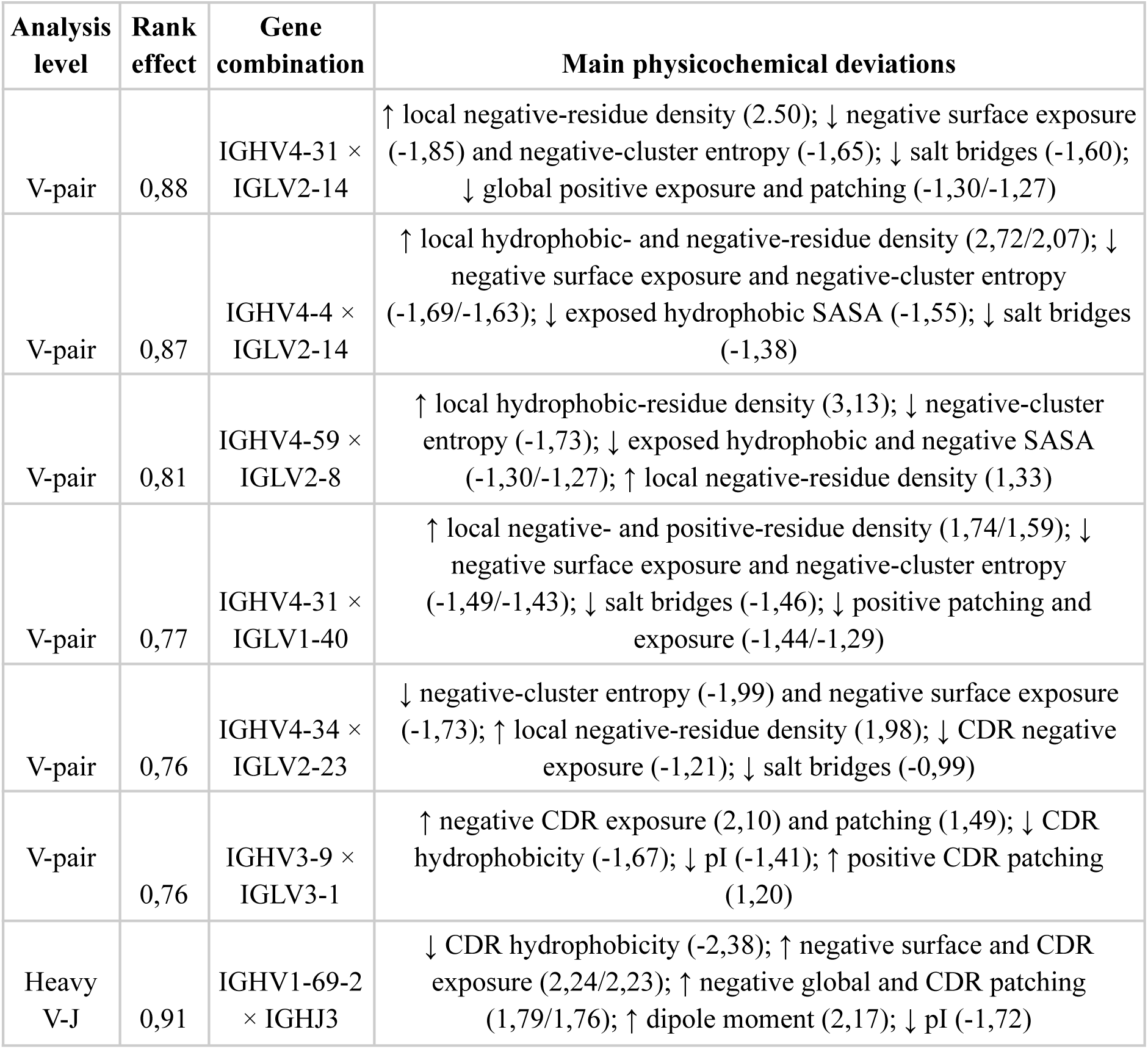
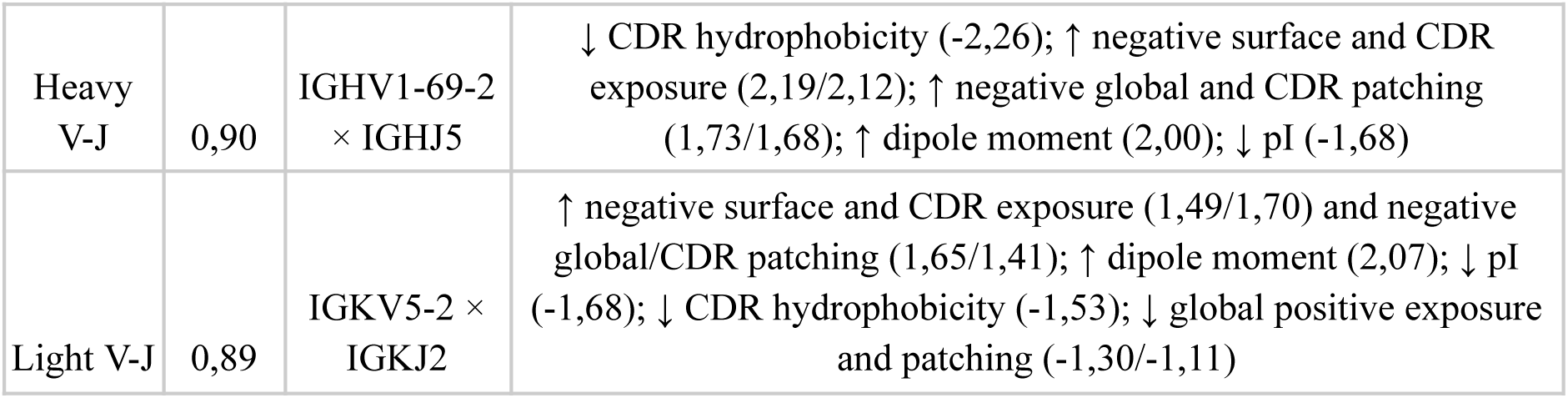
Germline gene combinations most strongly associated with greater distance from the therapeutic Fv profile.

| Analysis level | Rank effect | Gene combination | Main physicochemical deviations |
| --- | --- | --- | --- |
| V-pair | 0,88 | IGHV4-31 × IGLV2-14 | ↑ local negative-residue density (2,50); ↓ negative surface exposure (-1,85) and negative-cluster entropy (-1,65); ↓ salt bridges (-1,60); ↓ global positive exposure and patching (-1,30/-1,27) |
| V-pair | 0,87 | IGHV4-4 × IGLV2-14 | ↑ local hydrophobic- and negative-residue density (2,72/2,07); ↓ negative surface exposure and negative-cluster entropy (-1,69/-1,63); ↓ exposed hydrophobic SASA (-1,55); ↓ salt bridges (-1,38) |
| V-pair | 0,81 | IGHV4-59 × IGLV2-8 | ↑ local hydrophobic-residue density (3,13); ↓ negative-cluster entropy (-1,73); ↓ exposed hydrophobic and negative SASA (-1,30/-1,27); ↑ local negative-residue density (1,33) |
| V-pair | 0,77 | IGHV4-31 × IGLV1-40 | ↑ local negative- and positive-residue density (1,74/1,59); ↓ negative surface exposure and negative-cluster entropy (-1,49/-1,43); ↓ salt bridges (-1,46); ↓ positive patching and exposure (-1,44/-1,29) |
| V-pair | 0,76 | IGHV4-34 × IGLV2-23 | ↓ negative-cluster entropy (-1,99) and negative surface exposure (-1,73); ↑ local negative-residue density (1,98); ↓ CDR negative exposure (-1,21); ↓ salt bridges (-0,99) |
| V-pair | 0,76 | IGHV3-9 × IGLV3-1 | ↑ negative CDR exposure (2,10) and patching (1,49); ↓ CDR hydrophobicity (-1,67); ↓ pI (-1,41); ↑ positive CDR patching (1,20) |
| Heavy V-J | 0,91 | IGHV1-69-2 × IGHJ3 | ↓ CDR hydrophobicity (-2,38); ↑ negative surface and CDR exposure (2,24/2,23); ↑ negative global and CDR patching (1,79/1,76); ↑ dipole moment (2,17); ↓ pI (-1,72) |
| Heavy V-J | 0,90 | IGHV1-69-2 × IGHJ5 | ↓ CDR hydrophobicity (-2,26); ↑ negative surface and CDR exposure (2,19/2,12); ↑ negative global and CDR patching (1,73/1,68); ↑ dipole moment (2,00); ↓ pI (-1,68) |
| Light V-J | 0,89 | IGKV5-2 × IGKJ2 | ↑ negative surface and CDR exposure (1,49/1,70) and negative global/CDR patching (1,65/1,41); ↑ dipole moment (2,07); ↓ pI (-1,68); ↓ CDR hydrophobicity (-1,53); ↓ global positive exposure and patching (-1,30/-1,11) |

Among heavy-chain V–J combinations represented by at least 1,000 antibodies, the strongest associations with greater distance from the therapeutic profile were IGHV1-69-2 × IGHJ3 and IGHV1-69-2 × IGHJ5. Both showed nearly identical acidic, weakly hydrophobic, highly dipolar profiles with reduced aromatic surface exposure. Among light-chain V-J pairs, IGKV5-2 × IGKJ2 showed the strongest association with greater distance. Its profile combined increased negative charge locally and globally, reduced CDR hydrophobicity, and strongly reduced global positive exposed SASA and patch, while positive CDR patch magnitude remained nearly unchanged.

Although descriptor missingness due to minimization failure was modest overall, its enrichment among several germline genes (see Methods) should be considered when interpreting germline-specific effects.

We next tested whether divergence from inferred germline was associated with therapeutic-profile distance or individual physicochemical descriptors (Supplementary Section 1). No association met the predefined across-source robustness criteria. Within individual sources, however, greater framework divergence was associated with increased positive CDR exposure in nine sources (Spearman ρ = 0.25–0.34, BH-adjusted q ≤ 0.05), whereas positive CDR patching was not recurrent. Framework divergence was also associated with lower `scm_neg_ff19sb` values in three sources (ρ = -0.30 to -0.27).

## Discussion

Predictive performance varied with assay type and was generally higher for readouts closely related to static Fv physicochemistry. Hydrophobicity- and charge-dominated measurements were among the most consistently predicted across datasets, and assay group remained significantly associated with performance after accounting for dataset identity. Expanding the feature- and model-selection space did not improve mean performance. In the low-n regime of current public developability datasets, additional modeling flexibility therefore did not compensate for weak, heterogeneous, or context-dependent assay signal.

Chromatographic hydrophobicity provided the clearest example of a comparatively direct physicochemical readout. Across independent HIC and SMAC datasets, models repeatedly selected exposed hydrophobicity, hydrophobic patches, and aromatic surface descriptors. By comparison, self-association and cross-interaction assays drew on varying combinations of charge, hydrophobicity, dipole moment, and surface organization, and their predictability differed substantially even among assays intended to probe related liabilities. One interpretation is that exposed hydrophobicity constitutes a relatively low-dimensional molecular axis: HIC retention is sufficiently close to the static surface chemistry represented by kitAb for a similar signal to recur across antibody panels, whereas self-association and nonspecific-interaction phenotypes may depend more strongly on context-dependent molecular determinants that are only partially represented by static Fv descriptors.

Some endpoints appeared to contain little information beyond relatively simple sequence-derived physicochemical properties: cIEF and heparin retention were largely captured by bulk electrostatic features. For several endpoints, sequence-only descriptors did not merely approach the structural representations but outperformed them. This was most pronounced for thermostability, where the sequence baseline exceeded kitAb for Jain 2024 Tm and Garbinski Tm1, and also exceeded full PROPERMAB for these endpoints and for Jain 2024 PSR. These results do not imply that structural information is intrinsically irrelevant, because plausible structural descriptors were repeatedly selected. Rather, in small datasets, expanding the representation introduces additional correlated and potentially noisy dimensions whose contributions must be estimated from limited observations, which may offset the additional information provided by structure. The Ginkgo analysis also highlighted the contribution of external antibody metadata. Removing subtype annotations reduced performance across all four representations, with particularly large effects for several thermostability endpoints. Because these annotations were supplied to every representation, feature access was controlled across methods, but the absolute performance achieved on Ginkgo cannot be attributed solely to the intrinsic descriptor sets.

The most informative distinction between natural and therapeutic antibodies was how physicochemical properties were organized across the Fv surface. Therapeutics had the greatest global positive exposed surface while showing much smaller positive CDR patches than those observed in some natural subsets. A similar pattern emerged for aromatic residues: therapeutics displayed high aromatic exposure without a corresponding increase in contiguous CDR hydrophobic patches, while aromatic-charge synergy was also elevated. Therapeutic antibody physicochemistry may therefore be distinguished partly by surface patterning rather than composition alone.

Therapeutic antibodies did not resemble a simple continuation of the physicochemical patterns observed across the naïve, mixed, memory, and ASC/plasma repertoire groups. Natural subsets showed coordinated changes in CDR charge organization and pI, whereas therapeutics departed from those patterns in several dimensions; for example, they maintained relatively small positive CDR patches despite high global positive exposure. They also combined shorter CDR3 regions with increased CDR proline content. Therapeutics therefore occupied a physicochemical state that was not a simple extension of the differences observed among natural repertoire groups. Unlike natural antibodies, they are subject to additional constraints in which antigen recognition must coexist with manufacturability. Their physicochemical profiles may therefore combine properties uncommon within any single natural repertoire group.

Therapeutic-profile proximity was not determined consistently by individual chain genes; instead, some of the strongest patterns emerged after considering heavy–light combinations. The combinations showing the strongest associations with greater therapeutic-profile similarity were all IGHV3 × IGKV1 pairings, while several of the strongest associations with greater distance involved IGHV4 together with lambda light chains.

The IGKV5-2 example illustrates why individual developability-related properties should not be conflated with overall therapeutic suitability. IGKV5-2 was strongly displaced from the multivariate therapeutic profile through an unusually acidic, negatively charged and highly dipolar phenotype, yet the literature notes that this germline has previously been associated with low polyreactivity. A property that is favorable along one developability axis may therefore still place an antibody away from the profile characteristic of therapeutics.

Individual geometry-sensitive descriptors varied appreciably among structure predictors, yet downstream predictive performance was similar across backends. kitAb also performed comparably to the more computationally intensive PROPERMAB representation on average while reducing descriptor-calculation time by roughly an order of magnitude in the single-structure timing benchmark. This suggests that atomistic predicted structures can provide a stable basis for deriving coarse-grained physicochemical descriptors, even when individual geometry-sensitive features vary between structure predictors. Such a level of representation is particularly appropriate for large-library screening, where robustness and scalability may be more valuable than atomistic detail whose additional information cannot be reliably learned from datasets containing only tens or hundreds of measurements.

Several limitations should be considered. First, kitAb characterizes the Fv, whereas many developability assays are also influenced by Fc/subclass, formulation composition, concentration, pH, and other experimental conditions. Second, public developability datasets are small. Random splits used for shared-parent series primarily estimate local interpolation within an engineered sequence neighborhood, and flat-CV results for endpoints with <80 samples do not provide the same protection against model-selection optimism as nested CV. Third, structural analyses used predicted rather than experimental Fv structures, and the repertoire atlas relied on ABodyBuilder2 alone; although downstream model performance was similar across structure backends, some individual descriptors were method-sensitive. Fourth, repertoire comparisons are affected by biological and technical heterogeneity in OAS and PairedAbNGS, by inferred germline assignments, and by nonrandom descriptor missingness for some germlines; they should therefore be interpreted as cross-sectional associations rather than maturation trajectories or causal effects. Finally, the 706 clinical-stage therapeutics constitute a selected reference population rather than a ground-truth definition of developability, and the runtime comparison was a single-structure benchmark.

The benchmark results point to different levels of physicochemical complexity across developability endpoints. Some readouts are dominated by relatively low-dimensional properties such as bulk charge or exposed hydrophobicity and can already be captured by simple representations. Others may depend more strongly on the spatial arrangement of those properties and benefit from structural descriptors. Still others plausibly depend on context-dependent intermolecular behavior that is not fully represented by descriptors of a static Fv. The therapeutic repertoire reinforces this distinction, showing that global surface composition can be retained while potentially unfavorable local organization is reduced.

## Conclusion

kitAb provides an open-source framework for scalable physicochemical characterization and assay-specific developability modeling of antibody Fv regions. Across 46 experimental endpoints, its interpretable descriptors performed similarly to PROPERMAB and exceeded TAP and the sequence-only baseline on average in sequence-aware or predefined evaluations. In the controlled single-structure CPU benchmark, kitAb descriptor calculation was approximately 11-fold faster than PROPERMAB. Predictive performance was strongly assay-dependent.

Applied to approximately 3.4 million paired natural antibodies, the same descriptor representation revealed systematic differences between natural repertoires and 706 Phase II or later therapeutic antibodies in CDR length, surface charge, hydrophobic and aromatic organization, and germline-associated physicochemical profiles. Together, these analyses position kitAb as a common physicochemical coordinate system: with labeled data, it supports assay-specific prediction; at repertoire scale, it enables comparison of antibodies within a multidimensional developability-relevant landscape. By combining computational efficiency, interpretability, and scalability, kitAb provides a practical framework for candidate prioritization and large-scale analysis of natural and therapeutic antibody space.

## Data and code availability

The kitAb source code, documentation, and analysis workflows are available at https://github.com/katyachemistry/kitAb/. Descriptor vectors for the paired natural-antibody dataset used in the repertoire-scale analyses are made available upon request. An interactive web server providing access to kitAb calculations and antibody physicochemical profiling is available at https://kitab-atlas.com/. All public developability datasets used for benchmarking are available from their original publications, repositories, or the FLAb2 benchmark repository at https://github.com/Graylab/FLAb [12].

