## Supplementary materials for "kitAb: an automated framework for assay-specific antibody developability modeling and large-scale landscape analysis"

### Section 1

#### Antibody structures

Antibody Fv structures were predicted with ABodyBuilder2 (ABB2), ABodyBuilder3 (ABB3) and FlashABB [20–22]. These three structure-prediction backends were used to assess cross-method variability in calculated descriptors, and the sensitivity of downstream machine-learning performance to the structure-generation method.

#### Structure processing

To assess within-method reproducibility, the complete structure processing procedure was repeated independently three times for each predicted Fv, and descriptors were calculated separately for the three resulting structures. Predicted Fv structures were refined using an ABodyBuilder2-adapted OpenMM protocol with OpenMM 8.4.0, PDBFixer 1.12, and the AMBER ff14SB force field. Missing heavy atoms, typically terminal OXT atoms, were added with PDBFixer. Hydrogens were subsequently added at pH 7 using OpenMM Modeller.addHydrogens. Because initial coordinates for newly added hydrogens contain a stochastic component, repeated preparation runs can begin from slightly different hydrogen configurations; subsequent restrained minimization can propagate these differences to nearby heavy-atom and side-chain geometries.

Structures were minimized in vacuo on the OpenMM CPU platform using L-BFGS energy minimization with harmonic positional restraints on N, CA, C, and CB atoms. Harmonic restraints were applied to backbone atoms during minimization and adaptively relaxed to maintain acceptable peptide-bond geometry. Non-proline cis peptide bonds were corrected using additional torsional restraints with progressively increased force constants. Refined structures were then checked for peptide-bond geometry, trans peptide configuration, L-stereochemistry at Cα, heavy-atom clashes, and strained side-chain bonds; problematic side chains were rebuilt and re-minimized when required. The refinement procedure was retried up to three times if validation criteria were not satisfied.

#### External software and parallelization

PROPKA 3.5.1 [26] was used to estimate pKa values for titratable residues and termini for charge-related descriptors. Absolute and relative solvent accessible surface area (SASA) values were computed with FreeSASA 2.1.2 [23] using the Shrake–Rupley algorithm and Ala–X–Ala tripeptide as a reference for relative SASA. Secondary structure and hydrogen-bond energies were obtained with DSSP 4.6.1 (mkdssp) [25]. Per-structure descriptor calculations were parallelized with GNU Parallel [54], and AutoML cross-validation jobs were distributed across CPU cores using Python multiprocessing.

#### Descriptors

Let $\mathcal{R}$ denote the set of all residues in the antibody Fv, and let ($\mathcal{C} \subseteq \mathcal{R}$) denote the strict IMGT-defined CDR residue set. The CDR-vicinity set, $\mathcal{V}$, comprised all CDR residues and all additional residues having at least one heavy atom within 5.0 Å of any CDR heavy atom:

$$\mathcal{V} = \mathcal{C} \cup r \in \mathcal{R} \setminus \mathcal{C} : \min_{c \in \mathcal{C}} d_{\mathrm{HA}}(r, c) \leq 5.0 \text{ Å}$$

where $d_{\mathrm{HA}}(r;\ c)$ denotes the minimum distance between any heavy atom of residue $r$ and any heavy atom of CDR residue $c$.

Four physicochemical residue classes were considered:

$$\mathcal{G} = \mathrm{neg}, \mathrm{pos}, \mathrm{aro}, \mathrm{hyd}$$

The corresponding residue-class sets were defined as follows:

$$\mathcal{R}_{\mathrm{neg}} = r \in \mathcal{R} : q(r) < -0.05$$

$$\mathcal{R}_{\mathrm{pos}} = r \in \mathcal{R} : q(r) > 0.05$$

$$\mathcal{R}_{\mathrm{aro}} = r \in \mathcal{R} : \mathrm{aa}(r) \in \mathrm{Phe}, \mathrm{Tyr}, \mathrm{Trp}, \mathrm{His}$$

$$\mathcal{R}_{\mathrm{hyd}} = r \in \mathcal{R} : \mathrm{aa}(r) \in \mathrm{Val}, \mathrm{Leu}, \mathrm{Ile}, \mathrm{Met}, \mathrm{Phe}, \mathrm{Trp}, \mathrm{Tyr}, \mathrm{Cys}$$

For each physicochemical class $g \in \mathcal{G}$, the corresponding CDR-vicinity subset was defined as:

$$\mathcal{V}_g = \mathcal{V} \cap \mathcal{R}_g, \qquad g \in \mathcal{G}$$

Here, $q(r)$ denotes the PROPKA-derived fractional charge of residue $r$ at pH 7.5, and $\mathrm{aa}(r)$ denotes its amino acid identity. The negatively charged class consisted predominantly of Asp and Glu, whereas the positively charged class primarily comprised Lys, Arg, and protonated His. Ala, Gly, and Pro were excluded from the hydrophobic class to focus on stronger patch-forming side chains.

##### SASA sums

SASA-based descriptors were computed from residue-level FreeSASA records. For each physicochemical class $g \in \mathcal{G}$, the total absolute side-chain SASA was defined as:

$$A_g^{\mathrm{all}} = \sum_{r \in \mathcal{R}_g} A_r^{\mathrm{side}}$$

where $A_r^{\mathrm{side}}$ is the absolute side-chain SASA of residue $r$, measured in Å².

Because exposure thresholds vary across studies, residues were classified as exposed using a 20% relative side-chain SASA cutoff. Let $s_r^{\mathrm{side}}$ denote the relative side-chain SASA of residue $r$. The exposed-residue set was defined as:

$$\mathcal{E} = r \in \mathcal{R} : s_r^{\text{side}} \geq 0.20$$

For each class, the exposed class-specific residue set was therefore:

$$\mathcal{R}_g^{\exp} = \mathcal{R}_g \cap \mathcal{E}$$

The total absolute SASA contributed by exposed residues of class *g* was calculated as:

$$A_g^{\exp} = \sum_{r \in \mathcal{R}_g^{\exp}} A_r^{\text{side}}$$

These calculations yielded `aro_all_sasa`, `aro_exposed_sasa`, `neg_all_sasa`, `neg_exposed_sasa`, `pos_all_sasa`, `pos_exposed_sasa`, `hyd_all_sasa`, and `hyd_exposed_sasa`.

Using the previously defined CDR-vicinity subset $\mathcal{V}_g = \mathcal{V} \cap \mathcal{R}_g$, its exposed subset was defined as:

$$\mathcal{V}_g^{\exp} = \mathcal{V}_g \cap \mathcal{E}$$

The absolute side-chain SASA of exposed residues within the CDR vicinity was then calculated as:

$$A_g^{\text{vic}} = \sum_{r \in \mathcal{V}_g^{\exp}} A_r^{\text{side}}$$

These descriptors were exported as `aro_sasa_cdr`, `neg_sasa_cdr`, and `pos_sasa_cdr`.

CDR-vicinity exposure descriptors were calculated by summing the relative side-chain SASA values of the same exposed residues:

$$E_g^{\text{vic}} = \sum_{r \in \mathcal{V}_g^{\exp}} s_r^{\text{side}}$$

These descriptors were exported as `aro_exposure_cdr`, `hyd_exposure_cdr`, `neg_exposure_cdr`, and `pos_exposure_cdr`.

#### Hydrophobicity scores

Residue hydrophobicity was assigned according to the Wimley-White interfacial scale, with unrecognized residues assigned a value of zero. Let *H*(*r*) denote the hydrophobicity of residue *r*. The overall hydrophobic score was calculated as:

$$\text{hyd}_{\text{score}} = \sum_{r \in \mathcal{R}} H(r)$$

CDR-vicinity hydrophobicity was calculated as the mean exposure-weighted hydrophobicity across residues in the CDR-vicinity set:

$$H_{\text{CDR}}^{\text{exp}} = \frac{1}{|\mathcal{V}|} \sum_{r \in \mathcal{V}} H(r),\, s_r^{\text{side}}$$

where $s_r^{\text{side}}$ is the relative side-chain SASA of residue $r$. Exposed hydrophobic side chains therefore contribute more strongly than buried residues. The descriptor was exported as `exposure_weighted_hyd_score_cdr`.

#### Local geometric flatness

Local planarity descriptors (`neg_planarity_cdr`, `pos_planarity_cdr`, `aro_planarity_cdr`, and `hyd_planarity_cdr`) were computed for exposed CDR-vicinity residues from selected physicochemical classes.

For each atom, $n$ neighboring heavy atoms within 8 Å were identified. Their Cartesian coordinates were then arranged into a matrix $X \in \mathbb{R}^{n \times 3}$, with one row per atom. The coordinate matrix was centered by subtracting the neighborhood centroid $\bar{\mathbf{r}}$ from each row:

$$X_c = X - \bar{\mathbf{r}}$$

The covariance matrix was then calculated as:

$$C = \frac{1}{n} X_c^\top X_c$$

Eigenvalue decomposition of $C$ yielded eigenvalues ordered as $\lambda_1 \leq \lambda_2 \leq \lambda_3$. The atom-level planarity score was defined as:

$$P = \frac{\lambda_1}{\lambda_1 + \lambda_2 + \lambda_3}$$

Lower values indicate that the local atomic neighborhood is compressed along at least one spatial direction and is therefore less three-dimensionally isotropic. Such neighborhoods are consistent with locally planar or flattened arrangements. However, the score does not uniquely distinguish planar neighborhoods from other dimensionally collapsed configurations, such as nearly collinear atomic arrangements. Thus, “planarity” is used here as an operational description of local geometric flattening rather than as a strict classification of a neighborhood as planar.

Residue-level scores were calculated as the mean atom-level score across the heavy atoms of each residue. Atoms whose 8 Å neighborhood contained fewer than four heavy atoms, including the central atom, were excluded. For each physicochemical class, residue-level scores were summed over exposed CDR-vicinity residues belonging to that class and normalized by the mean score across all exposed CDR-vicinity residues.

#### Hydrogen-bond descriptors

The two descriptors used different hydrogen-bond definitions. `hbond_density_cdr` was based on a custom geometry-based detector that included backbone and side-chain interactions, whereas `hbond_energy_density` used only backbone hydrogen bonds reported by DSSP.

Putative hydrogen bonds were identified using a heavy-atom geometric criterion derived from established protein H-bond detection approaches. Donor and acceptor atoms were assigned according to residue chemistry, and candidate pairs were required to have a donor-acceptor distance of at most 3.2 Å and a donor-antecedent-donor-acceptor angle of at least 120°. Atom-specific occupancy limits were applied to avoid overassignment. Same-chain backbone contacts separated by less than 3 residues were excluded. Residue contributions were weighted by relative SASA, using main-chain SASA for backbone atoms and side-chain SASA for side-chain atoms:

$$w_r^{\mathrm{HB}} = \sum_{x \in \mathcal{A}_r^{\mathrm{HB}}} w_x$$

$$\mathrm{hbond\_density\_cdr} = \frac{1}{|\mathcal{V}|} \sum_{r \in \mathcal{V}} w_r^{\mathrm{HB}}$$

Here, $\mathcal{A}_r^{\mathrm{HB}}$ is the multiset of donor and acceptor atoms assigned to residue $r$ across retained interactions, and $w_x$ is the corresponding relative SASA.

Separately, `hbond_energy_density` was calculated from DSSP backbone hydrogen bonds. For each interaction with energy $E$, both residues received a main-chain-SASA-weighted contribution:

$$\Delta e_r = s_r^{\mathrm{main}} |E|, \qquad \Delta e_{r'} = s_{r'}^{\mathrm{main}} |E|$$

$$\mathrm{hbond\_energy\_density} = \frac{1}{|\mathcal{R}_{\mathrm{HB}}|} \sum r \in \mathcal{R}_{\mathrm{HB}} e_r$$

where $r$ and $r'$ are the bonded residues, $s_r^{\mathrm{main}}$ is relative main-chain SASA, and $\mathcal{R}_{\mathrm{HB}}$ contains residues with nonzero DSSP-derived contributions. Descriptors were set to zero when no qualifying interactions were found.

#### Patch area descriptors

Residues were clustered separately by class using DBSCAN on Cα coordinates, without restricting to exposed residues. To reflect the longer spatial range of electrostatic interactions and the more compact packing expected for hydrophobic patches, parameters were set to $\varepsilon = 10$ Å and min_samples $= 2$ for charged residues, and $\varepsilon = 5$ Å and min_samples $= 3$ for hydrophobic residues. Residues classified as noise were excluded.

For each physicochemical class $g \in \mathcal{G}$, let $\mathcal{R}g^{\mathrm{clust}} \subseteq \mathcal{R}g$ denote the residues assigned by DBSCAN to clusters. Patch area was then calculated as:

$$A_g^{\text{patch}} = \sum_{r \in \mathcal{R}g^{\text{clust}}} Ar^{\text{side}}$$

where $A_r^{\text{side}}$ is the absolute side-chain SASA of residue $r$.

The same calculation was applied to the full Fv and CDR-vicinity residues, yielding `pos_patch`, `neg_patch`, `hyd_patch`, `pos_patch_cdr`, `neg_patch_cdr`, and `hyd_patch_cdr`.

#### Cluster entropy

For each charged class $g \in \text{neg}, \text{pos}$, the side-chain SASA of each cluster was expressed as a fraction $p_{g,k}$ of the total clustered side-chain SASA for that class. Cluster entropy was then calculated as:

$$H_g = -\sum_k p_{g,k} \log p_{g,k}$$

Higher values indicate that charged surface area is distributed across several similarly sized clusters, whereas lower values indicate one or a few dominant clusters. The descriptors were exported as `neg_cluster_entropy` and `pos_cluster_entropy`.

#### Pair-correlation enrichment descriptors

Exposed residues were grouped into negative, positive, and hydrophobic classes. For each class $g \in \text{neg}, \text{pos}, \text{hyd}$, let $\mathcal{E}g = \mathcal{E} \cap \mathcal{R}g$ denote the exposed residues belonging to that class.

For every unordered pair of distinct residues $r_i, r_j \subset \mathcal{E}g$, the distance between their Cα atoms was calculated. A pair was assigned to a shell only when the distance between the two Cα atoms fell within that interval: $4 \leq d_{\text{C}\alpha}(r_i, r_j) < 7\ \text{Å}$ or $7 \leq d_{\text{C}\alpha}(r_i, r_j) < 10\ \text{Å}$. For each class $g$ and distance shell $b$, the observed number of same-class residue pairs, $N_{g,b}^{\text{obs}}$, was compared with a random-surface null distribution. In each randomization, $|\mathcal{E}_g|$ residues were sampled from all exposed residues, and the number of pairs falling within the same distance shell was counted. The normalized pair enrichment was defined as:

$$\text{PCF}g, b = \frac{Ng, b^{\text{obs}}}{M^{-1} \sum_{m=1}^{M} N_{g,b}^{(m)}}$$

where $N_{g,b}^{(m)}$ is the pair count in randomization $m$ and $M$ is the number of randomizations. For each residue class and distance shell, 3000 randomizations were performed using a fixed random seed of 42. Values above $1$ indicate enrichment of same-class pairs relative to random placement on the exposed surface, whereas values below $1$ indicate depletion.

The resulting descriptors were `pcf_neg_4w3A`, `pcf_neg_7w3A`, `pcf_pos_4w3A`, `pcf_pos_7w3A`, `pcf_hyd_4w3A`, and `pcf_hyd_7w3A`. A value of $0.0$ was returned when there were fewer than two residues in the selected class.

#### SCM- and SAP-like scores

For the SCM-like descriptors, the local electrostatic environment of each atom was estimated by summing the ff19SB partial charges of all other atoms within $10.0\ \text{Å}$. Let $Q_y^{\text{ff19SB}}$ denote the partial charge of atom $y$ and $\mathcal{B}_{10}(x)$ the neighborhood of atom $x$:

$$\text{SCM}x = \sum_{y \in \mathcal{B}_{10}(x), y \neq x} Q_y^{\text{ff19SB}}$$

Negative and positive local charge environments were then accumulated separately:

$$S_{\text{SCM}}^{-} = \sum_{x:\text{SCM}x<0} \text{SCM}x, \qquad S_{\text{SCM}}^{+} = \sum_{x:\text{SCM}x>0} \text{SCM}x$$

These quantities were exported as `scm_neg_ff19sb` and `scm_pos_ff19sb`, respectively. The ff19SB partial charges were determined by the protonation states used during force-field parameterization and are distinct from the PROPKA-derived fractional residue charges used below.

SAP-like descriptors quantified exposure-weighted local aromatic and charge–aromatic environments across the Fv surface. For each residue *i*, let $s_i$ denote relative side-chain SASA and $q_i$ the PROPKA-derived fractional charge at pH $7.5$. Three exposure-weighted source terms were defined:

$$s_i^{+} = s_i \max(q_i, 0), \qquad s_i^{-} = s_i \min(q_i, 0), \qquad s_i^{\text{aro}} = s_i \mathbf{1}_{\{i \in \mathcal{R}_{\text{aro}}\}}$$

For each residue, $\mathcal{N}(i)$ contained all other residues having at least one heavy atom within $10.0\ \text{Å}$. Local positive, negative, and aromatic environments were calculated as:

$$P_i = \sum_{j \in \mathcal{N}(i)} s_j^{+}, \qquad N_i = \sum_{j \in \mathcal{N}(i)} s_j^{-}, \qquad A_i = \sum_{j \in \mathcal{N}(i)} s_j^{\text{aro}}$$

The final descriptors were:

$$S_{\text{aro}} = \sum_{i \in \mathcal{R}} s_i A_i, \qquad S_{\text{pos-aro}} = \sum_{i \in \mathcal{R}} s_i P_i A_i, \qquad S_{\text{aro-neg}} = \sum_{i \in \mathcal{R}} s_i A_i N_i$$

These were exported as `sap_aro_score`, `sap_pos_aro_synergy`, and `sap_aro_neg_synergy`.

#### Other descriptors

Unless stated otherwise, charge-based descriptors used PROPKA-derived fractional charges at the default pH values equal 7.5. Full-Fv `net_charge_by_pH` was computed at pH 3.0 and 7.5 from ionizable residues and chain termini:

$$Q(\text{pH}) = \sum_{r \in \mathcal{R}_{\text{ion}}} q_r(\text{pH}) + q_{\text{Nterm}}(\text{pH}) + q_{\text{Cterm}}(\text{pH})$$
.

The isoelectric point (`protein_pi`) was estimated as the pH in the range [0,14] where $Q(\text{pH}) = 0$. Heavy–light `charge_symmetry` was defined as:

$$\text{charge\_symmetry} = Q_H(\text{pH}) \times Q_L(\text{pH})$$
.

The dipole descriptor was computed from ff19SB atom partial charges after centering coordinates on the atomic centroid:

$$\boldsymbol{\mu} = \sum_i q_i(\mathbf{r}_i - \mathbf{r}_0), \qquad \|\boldsymbol{\mu}\| = \sqrt{\mu_x^2 + \mu_y^2 + \mu_z^2}$$
.

Inter-chain buried SASA was estimated as:

$$\text{inter\_chain\_buried\_sasa} = \text{SASA}_H + \text{SASA}_L - \text{SASA}_{HL}.$$

CDR composition descriptors included six fractions, each defined by formula:

$$f_g^{\text{CDR}} = \frac{|\mathcal{C} \cap \mathcal{R}_g|}{|\mathcal{C}|}$$
,

where classes were glycine, proline, aromatic residues, positively charged residues, negatively charged residues, and Gln/Asn; $\mathcal{C}$ is a set of CDR residues. Asp-Asp, Asp-Glu, and Tyr-Tyr motifs were counted in CDR-vicinity sequence fragments. `number_of_salt_bridges` was counted as unique oppositely charged residue pairs with at least one atom-atom contact within 4.0 Å. Each residue pair was counted once.

### AutoML

When predefined folds were unavailable, concatenated heavy- and light-chain Fv sequences were clustered with MMseqs2. Identity thresholds from 0.50 to 0.80 were tested, with five-fold and then four-fold assignments accepted only if all folds were non-empty and the largest fold contained no more than twice as many antibodies as the smallest. If no acceptable sequence-aware split was obtained, or if random splitting was explicitly specified, shuffled five-fold CV was used with three seeds.

All preprocessing and feature selection were performed within the relevant training partition. Features with relative standard deviation <0.01 were removed. For pairs with |Pearson's r| > 0.9, the feature showing the weaker absolute Pearson correlation with the target in the training set was discarded. Scaling, filtering, correlation pruning, SFS, and feature–target associations were therefore recomputed independently within each training fold.

For correlation-pruned SVR, all remaining features were used with a linear-kernel SVR. The two SFS strategies used either linear-kernel SVR or k-NN to guide forward selection, with improvement <0.02 as the stopping criterion and the number of selected features capped at 15% of the training-set size;

the final regressor was selected among linear regression, random forest, linear-kernel SVR, and k-NN. Elastic-net models used standardization, with regularization strength and L1 ratio selected by inner CV.

For endpoints with ≥80 samples, AutoML was evaluated by a nested CV. Within each outer training partition, the four modelling strategies and their internal settings were compared using pooled inner-validation Spearman correlation. The selected configuration was refitted on the complete outer training partition and used to predict the held-out outer fold. Outer-fold predictions were pooled to calculate Spearman correlation, Pearson correlation, $R^2$, and mean squared error. For smaller endpoints, the four strategies were evaluated by flat CV without an additional model-selection layer.

Feature recurrence was assessed using the mean pairwise Jaccard agreement between feature sets selected across outer folds for the modelling strategy chosen by inner-CV performance:

$$J_{ij} = \frac{|S_i \cap S_j|}{|S_i \cup S_j|}$$

Feature sets were defined as all features remaining after correlation pruning for correlation-pruned SVR, the SFS-selected features for SFS-based strategies, and features with nonzero coefficients for ElasticNet. Jaccard scores were averaged across all fold pairs and then across structure-model runs. Values range from 0 to 1, with higher values indicating greater feature recurrence across folds.

### Analysis

#### Descriptor variability

Descriptor sensitivity was evaluated for antibodies predicted with ABodyBuilder2, ABodyBuilder3, and FlashABB, each followed by three processing runs. Let $x_{ijmr}$ denote descriptor $j$ for antibody $i$, method $m$, and run $r$. Values were scaled by between-antibody variation:

$$s_j = \mathrm{SD}_i\left[\mathrm{median}_{m,r}(x_{ijmr})\right]$$

Run-to-run and between-method variability were calculated as:

$$V_{ijm}^{\mathrm{run}} = \frac{\max_r(x_{ijmr}) - \min_r(x_{ijmr})}{s_j}$$

$$V_{ij}^{\mathrm{method}} = \frac{\max_m(\bar{x}_{ijm}) - \min_m(\bar{x}_{ijm})}{s_j},$$

The medians of these values across antibodies were taken to describe the variability of each descriptor.

#### Predictions reproducibility

AutoML performance was compared across ABodyBuilder2, ABodyBuilder3, and FlashABB. For each dataset–endpoint combination, Spearman correlations were averaged across the three structure-prediction runs. The resulting matched endpoint-level correlations were compared across the three structure-generation methods using a Friedman test, with dataset–endpoint combinations treated as blocks. A Kruskal–Wallis test applied to the same method-specific correlation distributions was used as a secondary, unpaired sensitivity analysis.

Run-to-run variation was calculated as:

$$R_{a,m} = \frac{\max_r(\rho_{a,m,r}) - \min_r(\rho_{a,m,r})}{\text{mean}_r(\rho_{a,m,r})}$$

for assay *a*, method *m*, and run *r*.

Predictive performance was further analyzed using a linear mixed-effects model with Fisher-transformed Spearman correlation as the response. The model included log effective sample size, structure-generation method, assay group, split strategy, and CV nestedness as fixed effects, with dataset–target pair as a random intercept:

$$z(\rho) = \beta_0 + \beta_1 \log(N_{\text{eff}}) + \beta_2\text{StructureMethod} + \beta_3\text{AssayGroup} + \beta_4\text{SplitStrategy} + \beta_5\text{CVNestedness} + u_{\text{dataset-target}} + \varepsilon.$$

Fixed effects were tested by likelihood-ratio comparisons with models excluding the corresponding term. A sensitivity model additionally included dataset identity as a fixed effect.

#### Assessment of descriptor missingness

Structure preparation and minimization failed for 167,446 of 3,605,625 antibodies (4.64%). Missingness was non-uniform across several germline genes. It was enriched among IGHV4 antibodies (6.45%; OR = 1.61), particularly IGHV4-38-2 (14.69%; OR = 3.67), and among IGHJ6 antibodies (7.33%; OR = 1.94). Light-chain associations were weaker, including IGLV4-60 (10.8%; OR = 2.50), IGLV3-19 (6.74%; OR = 1.50), and IGKV3-20 (5.54%; OR = 1.23).

#### Distance-to-therapeutics germline genes enrichment

The 59-descriptor panel was reduced in two steps. Based on predefined methodological considerations, motif counts, 7 Å PCFs, total SASA, `charge_symmetry`, `hyd_score`, and most CDR composition fractions reduced the panel to 44 descriptors. Redundant descriptors were then removed when |Spearman ρ| > 0.8 in both the natural complete-case cohort and the 706 therapeutic antibodies, leaving 33 descriptors: `pcf_neg_4w3A`, `pcf_pos_4w3A`, `pcf_hyd_4w3A`, `neg_cluster_entropy`, `pos_cluster_entropy`, `pos_patch`, `neg_patch`, `pos_patch_cdr`, `neg_patch_cdr`, `hyd_patch_cdr`, `aro_exposed_sasa`, `neg_exposed_sasa`, `pos_exposed_sasa`, `hyd_exposed_sasa`, `hyd_exposure_cdr`, `neg_exposure_cdr`, `pos_exposure_cdr`, `exposure_weighted_hyd_score_cdr`, `hbond_density_cdr`, `scm_neg_ff19sb`, `scm_pos_ff19sb`, `sap_pos_aro_synergy`, `sap_aro_neg_synergy`, `hbond_energy_density`, `inter_chain_buried_sasa`, `dipole_moment_ff19sb`, `net_charge_by_pH.3`, `protein_pi`, `cdr3_length`, `number_of_salt_bridges`, `fraction_gly_in_cdr`, `fraction_pro_in_cdr`, and `fraction_gln_asn_in_cdr`.

The therapeutic profile was defined from the 706 Phase II-or-later Thera-SAbDab antibodies. Each descriptor was centered on the therapeutic median and scaled by its therapeutic IQR; therapeutic-profile distance was calculated as the root mean square across the 33 scaled descriptors.

Germline associations were tested within each biological source by comparing antibodies assigned to a given germline category with all other antibodies using Wilcoxon rank contrasts, with evidence combined across sources using the van Elteren test. Effect size was expressed as rank-biserial $r=2U/(n_1 n_0)-1$, with positive values indicating greater distance from therapeutics. Associations were considered robust if they remained significant after BH correction $q \leq 0.05$, had an absolute pooled rank-biserial effect size $|r| > 0.20$, and showed no significant between-source heterogeneity Cochran's Q, $P_Q \geq 0.05$. Descriptor-level enrichment was assessed by defining the lowest and highest 5% of values for each descriptor within each biological source and testing germline enrichment across sources using Cochran–Mantel–Haenszel tests; effects with $|\log_2 OR| \geq 0.25$ were retained.

Framework non-germline percentage, averaged across heavy and light chains, was used as the primary somatic-hypermutation proxy. Within-source Spearman correlations were combined by Fisher-z meta-analysis with Cochran's Q. Cross-source associations were retained only when at least 80% of contributing biological sources showed the same direction of effect.

#### Associations with non-germline fraction

Within each B-cell source, Spearman correlations were calculated between non-germline fraction and both therapeutic-profile distance and individual descriptors. Framework and CDR non-germline fractions were defined as the mean heavy- and light-chain mutation percentages. Source-specific correlations were combined by fixed-effect Fisher-z meta-analysis, with Cochran's Q used to assess heterogeneity. Within-source associations were retained for further consideration if they passed BH-FDR correction ($q \leq 0.05$) and had $|\rho| \geq 0.05$, the latter serving only as a minimal filter against near-zero effects. Cross-source associations additionally had to satisfy these criteria at the meta-analysis level, show concordant direction in at least 80% of sources, and show no significant heterogeneity ($p \geq 0.05$).

### Section 2

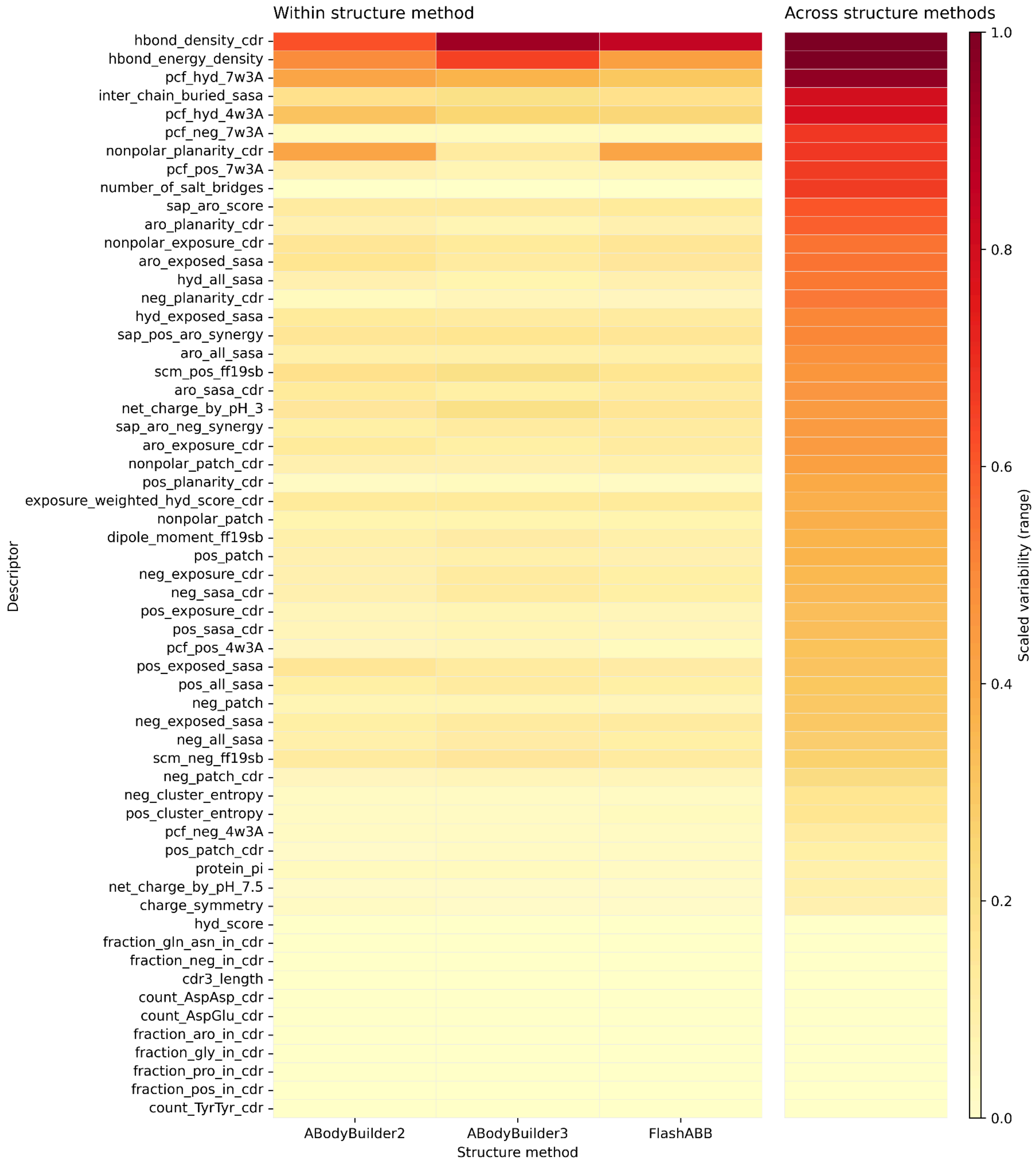


**Figure S1**. Heatmap of descriptor variability expressed as scaled range.

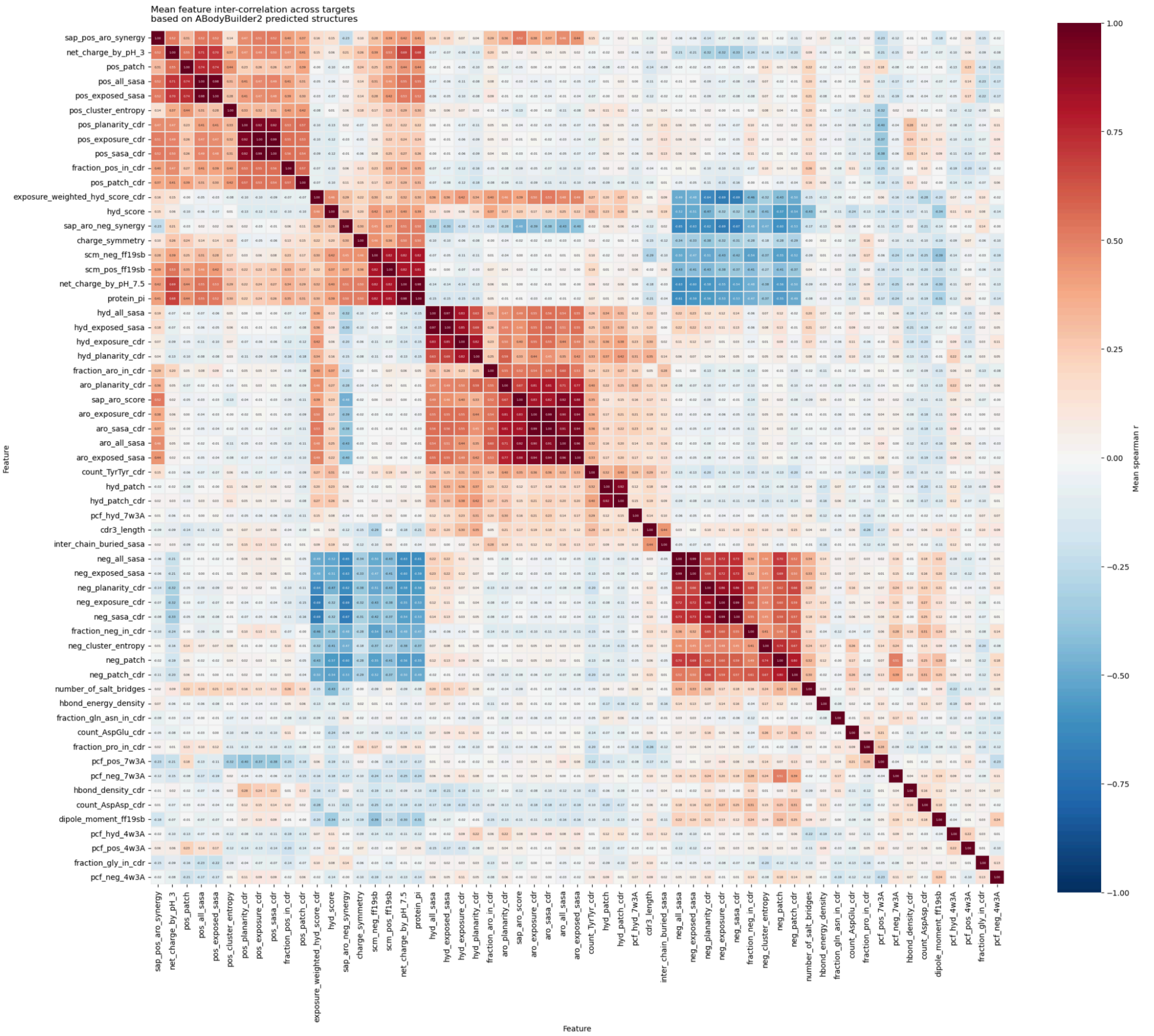


**Figure S2**. Mean Spearman correlations between developability descriptors, averaged across targets from 10 selected datasets (Table S1). Descriptors were calculated from ABodyBuilder2 predicted structures.

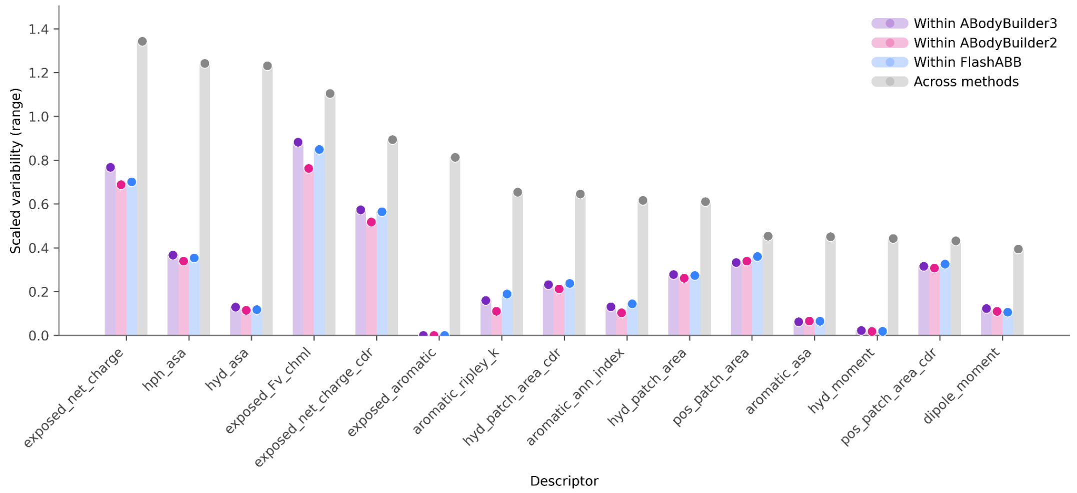


**Figure S3**. The 15 most variable PROPERMAB descriptors. Descriptor variability is expressed as the descriptor range, normalized by antibody-to-antibody variation (see Formula S1).

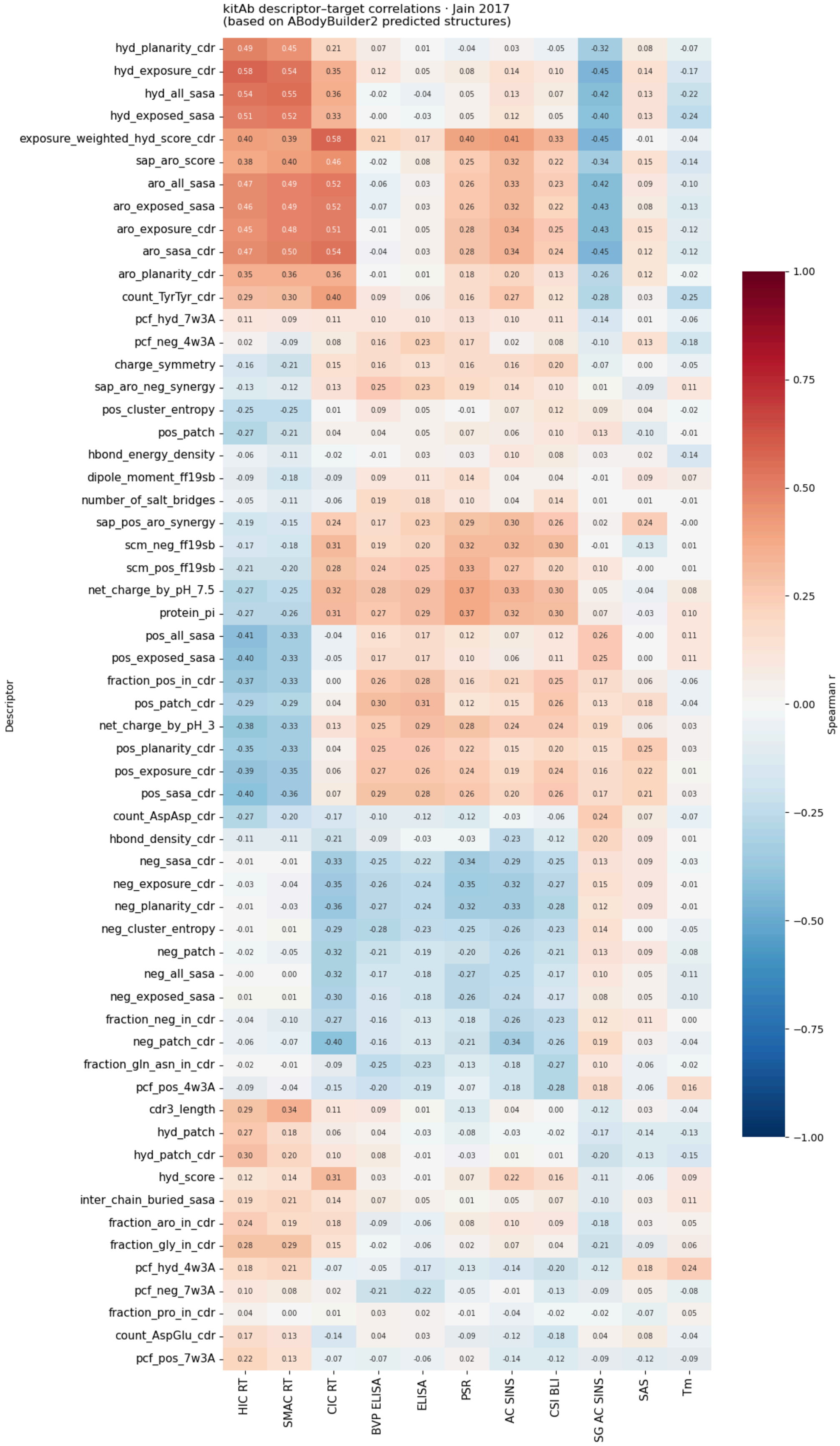


**Figure S4**. Heatmap of Spearman rank-correlation coefficients between assay endpoints and kitAb descriptors in the Jain 2017 dataset.

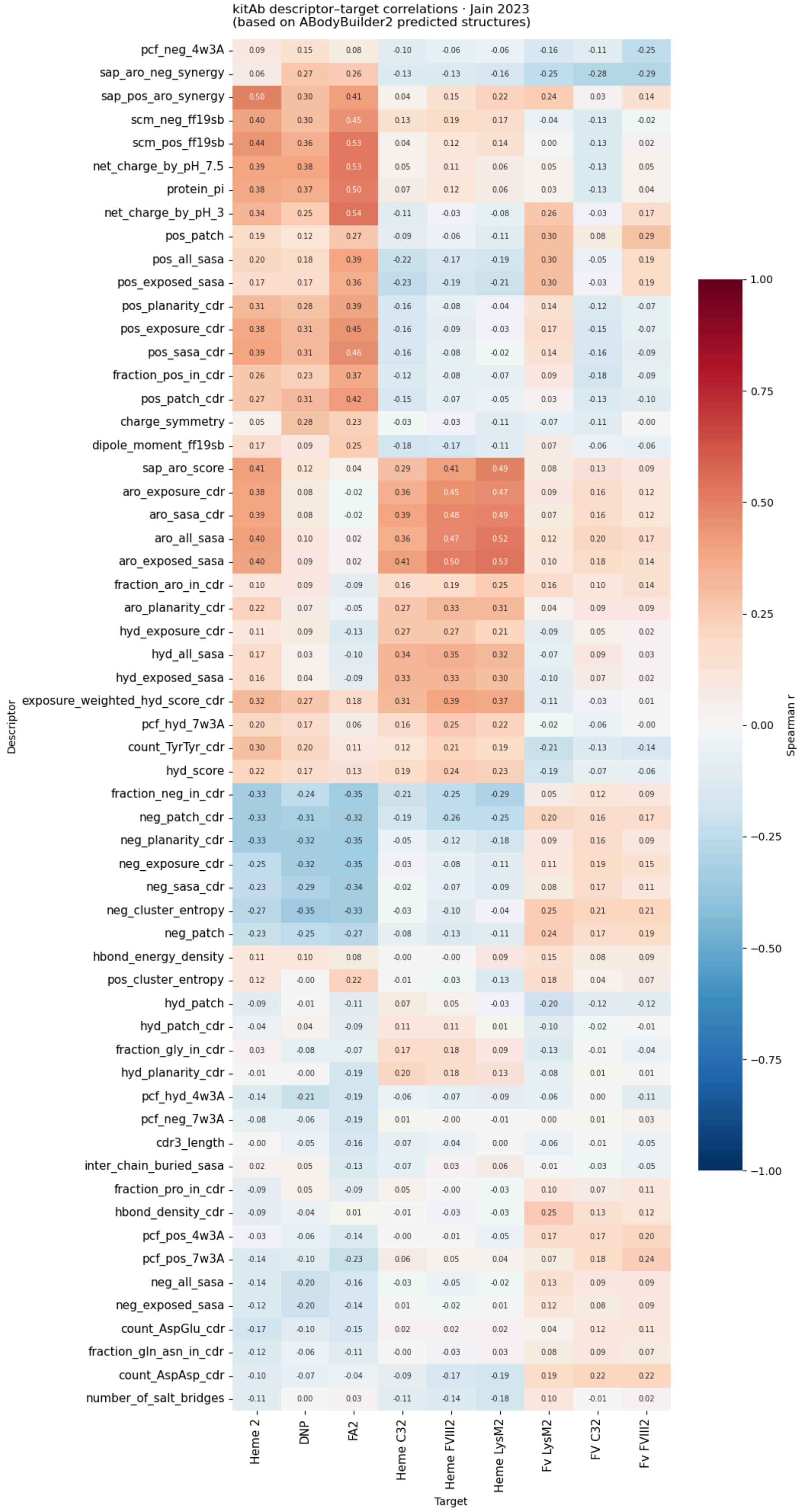


**Figure S5**. Heatmap of Spearman rank-correlation coefficients between assay endpoints and kitAb descriptors in the Jain 2023 dataset.

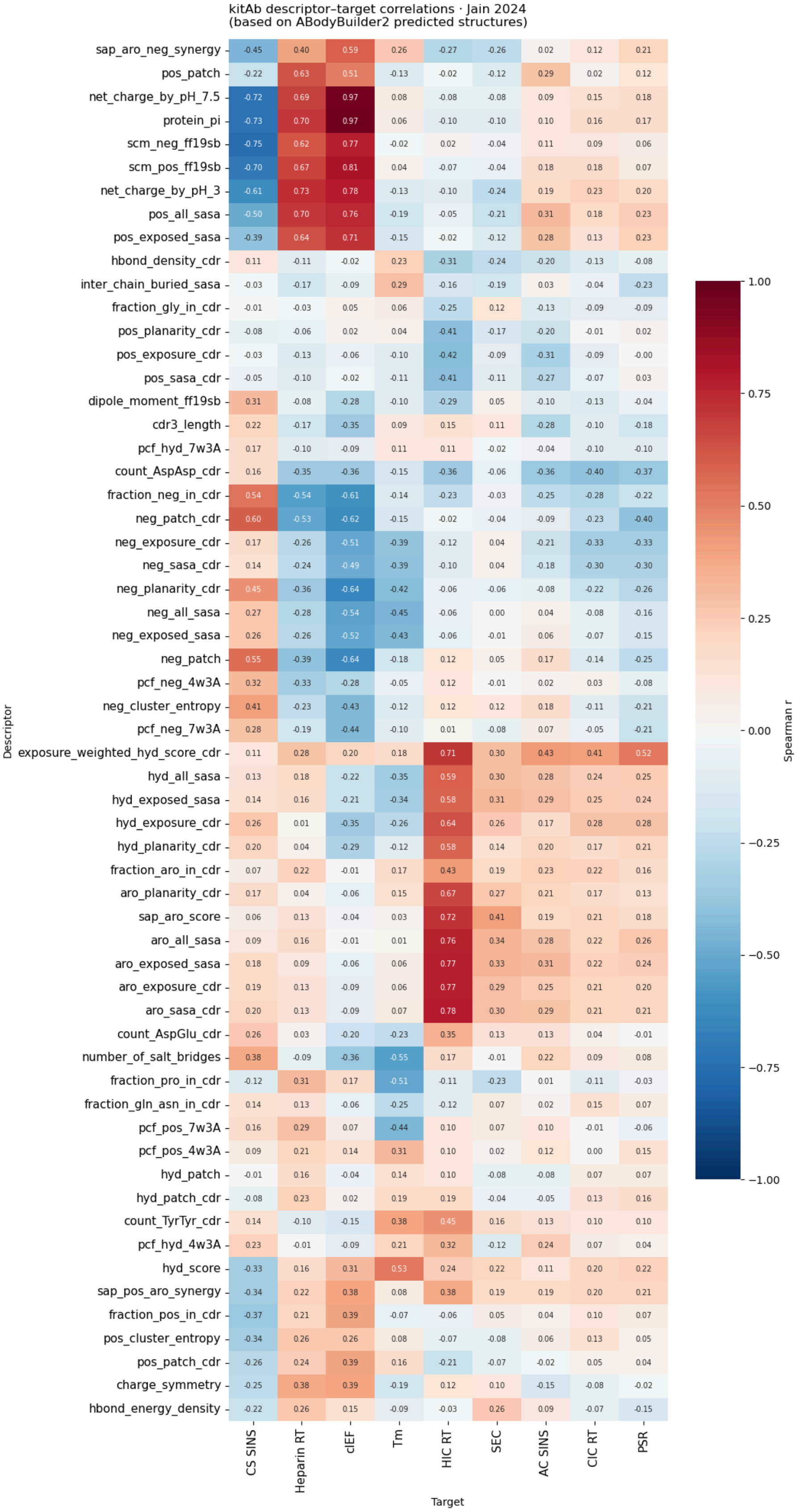


**Figure S6**. Heatmap of Spearman rank-correlation coefficients between assay endpoints and kitAb descriptors in the Jain 2024 dataset.

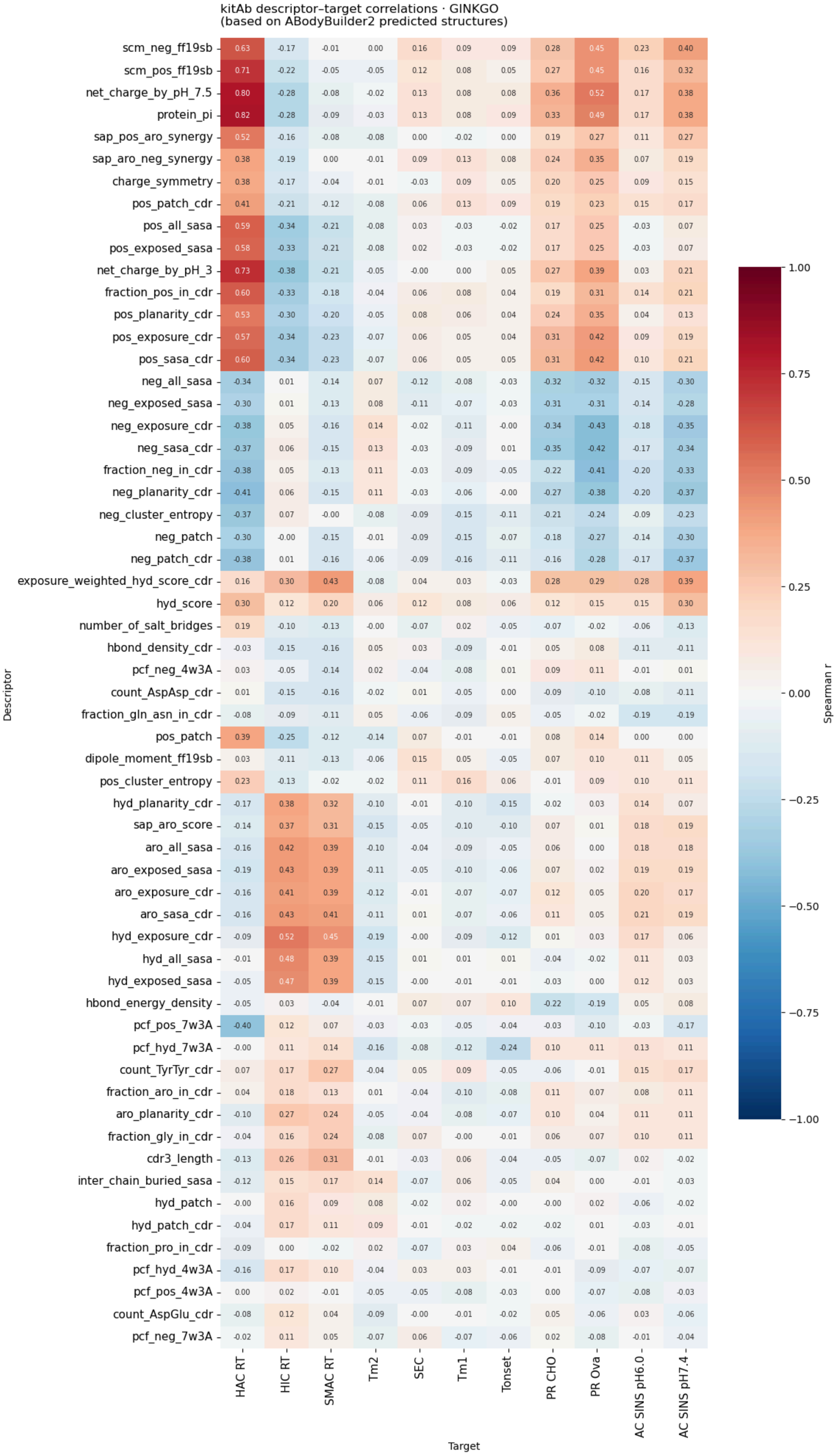


**Figure S7**. Heatmap of Spearman rank-correlation coefficients between assay endpoints and kitAb descriptors in the Ginkgo dataset.

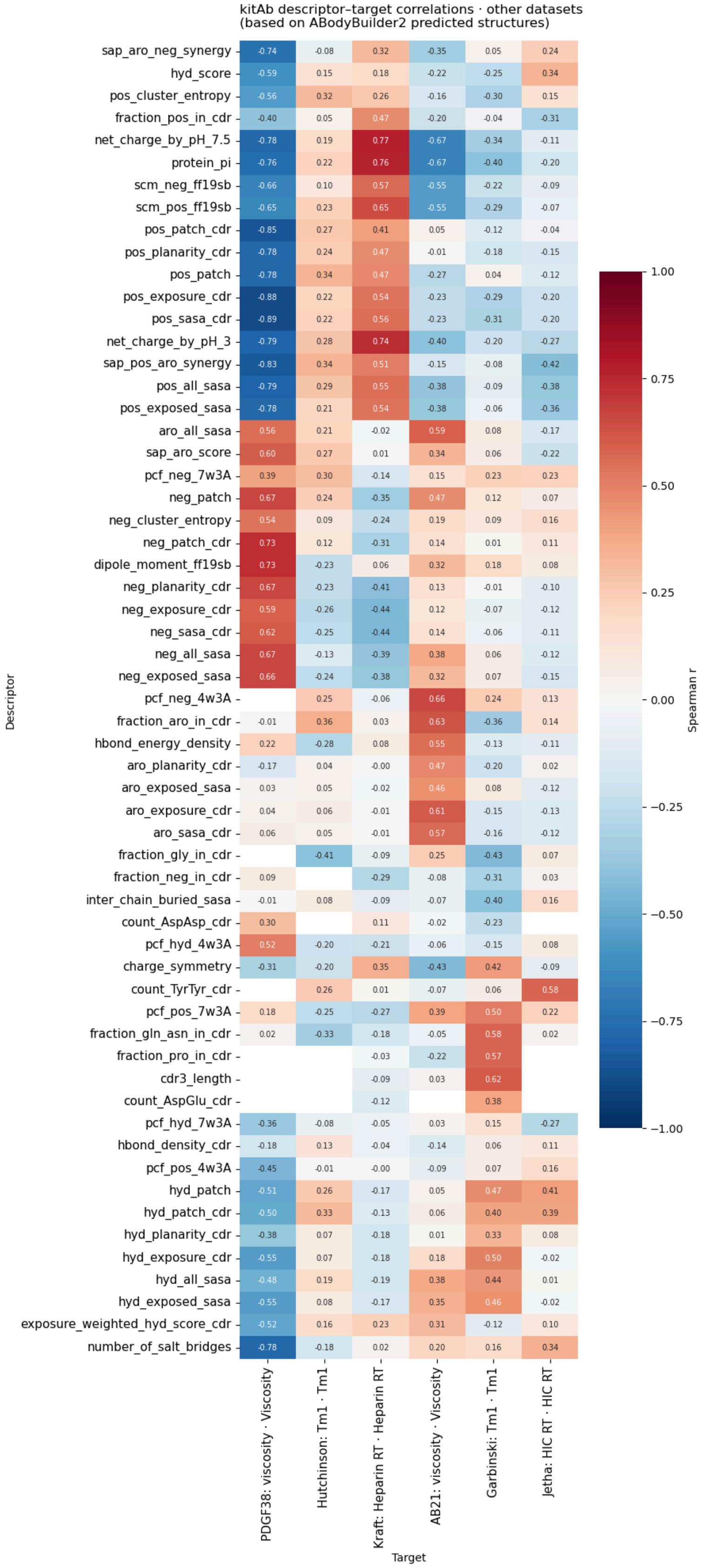


**Figure S8**. Heatmap of Spearman rank-correlation coefficients between assay endpoints and kitAb descriptors in the single-target datasets. Blank cells indicate descriptors that were constant within the corresponding dataset/endpoint.

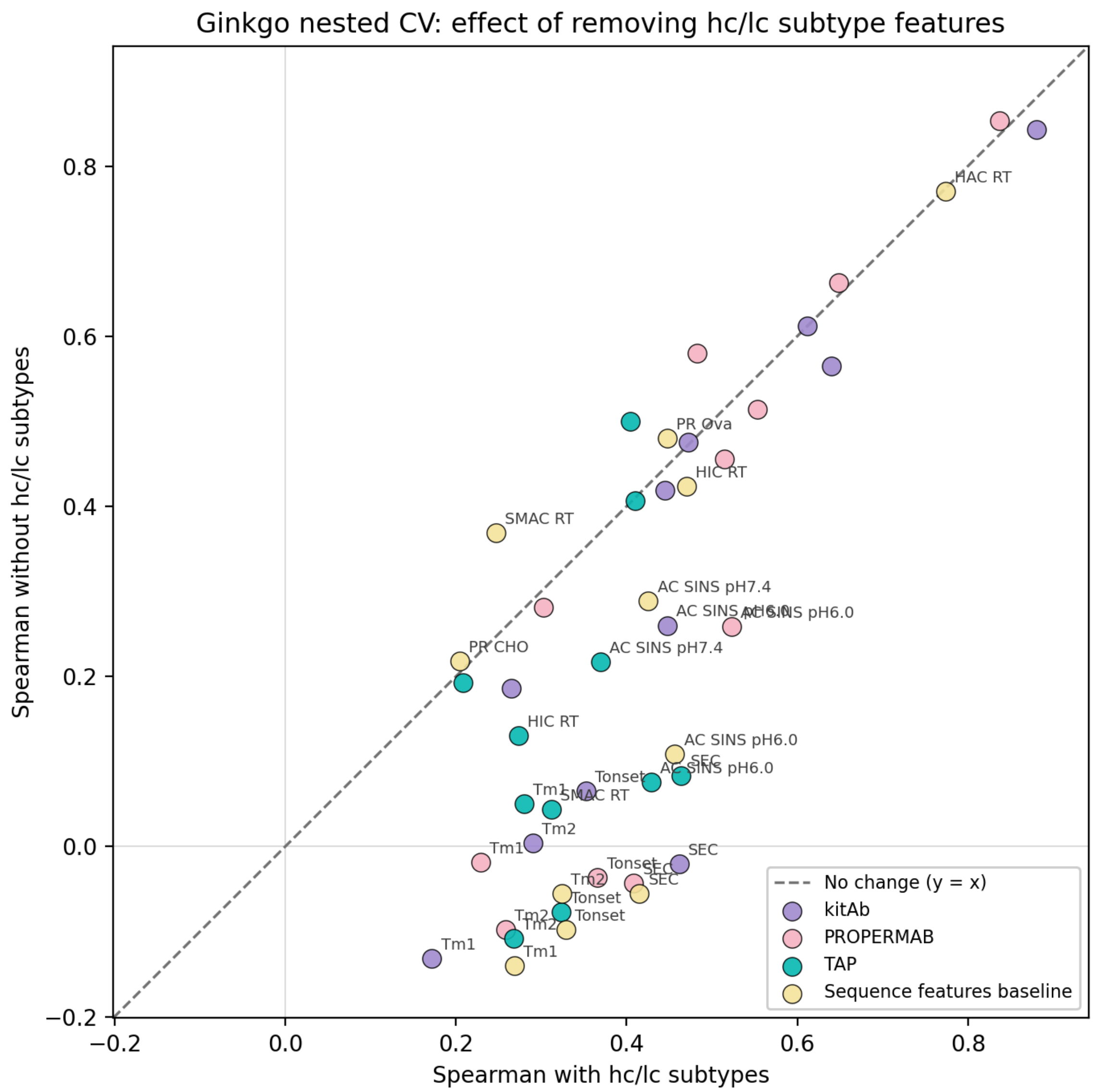


**Figure S9**. Spearman correlation of developability predictions on the Ginkgo dataset using ABodyBuilder2 structures, comparing the full feature sets with otherwise identical methods after removal of heavy and light chain class features.
